# Monkey-to-human transfer of brain-computer interface decoders

**DOI:** 10.1101/2022.11.12.515040

**Authors:** Fabio Rizzoglio, Ege Altan, Xuan Ma, Kevin L. Bodkin, Brian M. Dekleva, Sara A. Solla, Ann Kennedy, Lee E. Miller

## Abstract

Intracortical brain-computer interfaces (iBCIs) enable paralyzed persons to generate movement, but current methods require large amounts of both neural and movement-related data to be collected from the iBCI user for supervised decoder training. We hypothesized that the low-dimensional latent neural representations of motor behavior, known to be preserved across time, might also be preserved across individuals, and allow us to circumvent this problem. We trained a decoder to predict the electromyographic (EMG) activity for a “source” monkey from the latent signals of motor cortex. We then used Canonical Correlation Analysis to align the latent signals of a “target” monkey to those of the source. These decoders were as accurate across monkeys as they were across sessions for a given monkey. Remarkably, the same process with latent signals from a human participant with tetraplegia was within 90% of the with-monkey decoding across session accuracy. Our findings suggest that consistent representations of motor activity exist across animals and even species. Discovering this common representation is a crucial first step in designing iBCI decoders that perform well without large amounts of data and supervised subject-specific tuning.

## Introduction

Intracortical brain-computer interfaces (iBCIs) promise to restore voluntary movement to persons with paralyzed limbs. A kinematic iBCI uses a “decoder” to transform neural activity into signals that can be used to control a cursor or a robotic limb ^1–4^. By using the decoder, instead, to infer patterns of muscle activity (EMG), it is even possible to reanimate a user’s limb itself using Functional Electrical Stimulation (FES) to activate the paralyzed muscles ^5^. Decoders are typically computed with supervised learning methods, tuning parameters to minimize the error between decoder-predicted and motor output.

For a paralyzed human with no motor output, the standard approach is to use an “observation-based” decoder, computed to map M1 activity into an observed (and presumably attempted) kinematic trajectory ^6^. This approach is not feasible for a decoder that maps neural activity into EMGs, which cannot be observed. In this study, we sought to explore whether it would be possible to transfer a decoder trained with neural and EMG data recorded from a neurologically intact monkey to a human with tetraplegia.

At first glance, transferring a neural decoder across individuals seems impossible, as the inputs to the decoder are typically the activities of recorded neurons, which would obviously differ. To make this approach feasible, it would be necessary to find a common neural reference frame in which the representations of a given action were preserved across individuals.

Recent work has shown that the space of observed activity patterns of neurons in the primary motor cortex is constrained to a manifold of at most 10-20 dimensions ^7–10^. Thus, observed spiking of recorded neurons is effectively a noisy readout of a low-dimensional set of “latent signals” that correspond to the position of the neural activity within the manifold; the ever-changing set of neurons recorded over weeks and months means that our estimation of the manifold is embedded in different empirical neural spaces spanned by different coordinates ^11,12^. To achieve a stable representation of the neural manifold, we may perform a change of basis to align the manifold from a given day with that of another day. Canonical Correlation Analysis (CCA ^13^) provides an effective tool for aligning the latent signals, so as to make a fixed decoder effective across months or even years for the same monkey ^12^.

In this study, we examined the hypothesis that a similar alignment process might reveal consistent task representations not only across time, but also across individuals. We tested this hypothesis using data collected from three monkeys performing a task requiring production of forces in various directions about the wrist. We trained a decoder using neural and EMG data from one monkey and used it to make accurate EMG predictions from CCA-aligned latent neural signals from a second monkey. Critically, CCA alignment operates only on neural data, making it feasible to apply this strategy even to neural recordings from a paralyzed human. After developing and testing the methods with data from monkeys, we used the same methods to transfer a monkey-based decoder to a paralyzed human, using the aligned human neural activity to obtain estimates of the EMGs that would have been produced had the human not been paralyzed. These predicted EMG signals could be used to control an FES system providing voluntary control of the paralyzed muscles, as we have done previously in monkeys ^5^. Brain-controlled FES has been attempted in humans as well, but without the ability to infer attempted EMG directly ^14,15^. The same approach should be applicable to more complex, nonlinear decoders that require large amounts of training data, and are otherwise not feasible to use with humans.

## Results

### Aligned latent neural signals are very similar across monkeys

We trained three monkeys to perform an isometric wrist task that required them to exert forces to control a cursor on a computer monitor, moving it from a central target to one of eight peripheral targets (Fig. 1a, see Supplementary Methods). Each monkey was implanted with a 96-channel electrode array in the hand area of the primary motor cortex (M1), and with intramuscular leads in several forearm and hand muscles contralateral to the cortical implant. We recorded multi-unit spiking activity as well as electromyographic (EMG) activity as the monkeys performed the task. For each monkey, we recorded experimental data on two separate sessions, separated by two (monkey S), 29 (monkey J), or 46 (monkey K) days.

**Fig. 1.**
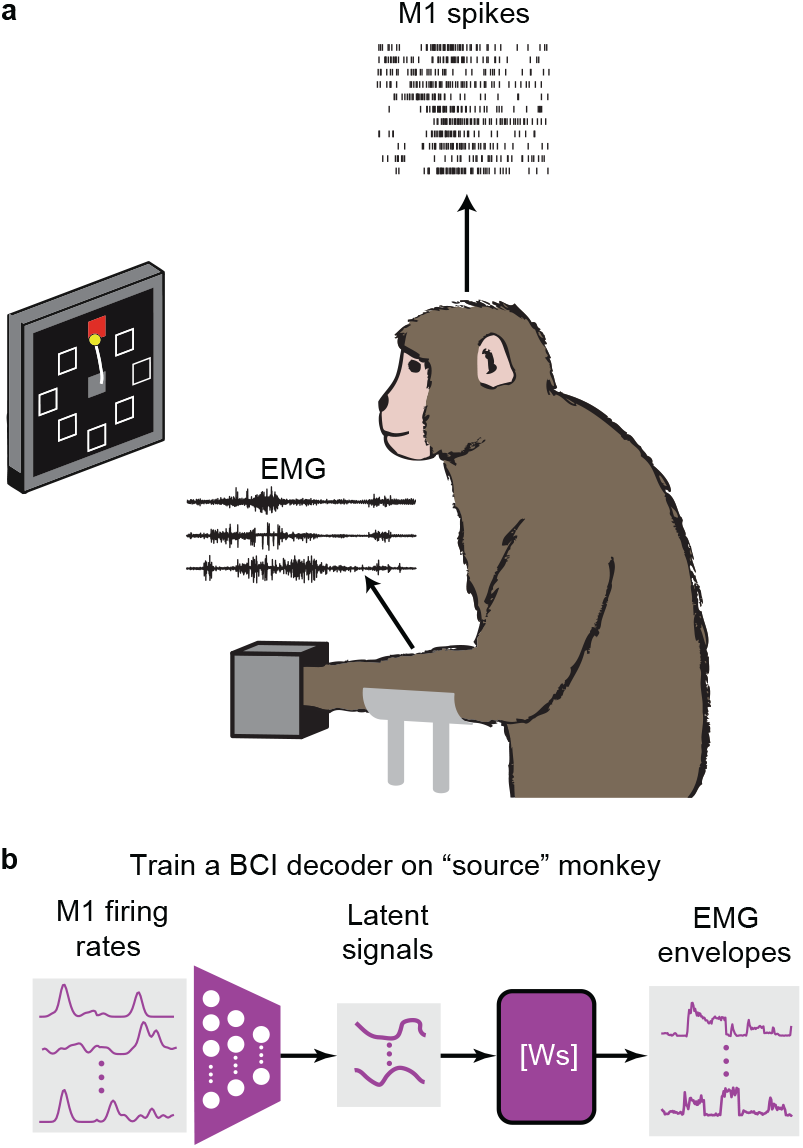
Training a fixed iBCI decoder. **a**, We record neural firing rates from M1 and EMGs from forearm and wrist muscles of a monkey (source monkey) trained to perform the isometric wrist task. **b**, The firing rates of the source monkey are projected in the low-dimensional neural manifold and the resulting latent signals are used as input to train the source-monkey iBCI decoder (Ws).

We used Parallel Analysis ^16,17^ to estimate an upper bound to the intrinsic dimensionality of the M1 data, which varied from 7 to 13 across datasets (see Methods). We applied Principal Component Analysis (PCA) to reduce the dimensionality of the neural data to the largest of these estimates. We used the resulting neural low-dimensional trajectories (termed *latent signals*) in training a Wiener cascade decoder ^18^ (see Methods) to predict the EMG signals of the *source* monkey (Fig. 1b).

We hypothesized that it would be possible to use this source-monkey decoder to make accurate EMG predictions using the latent signals from a second monkey (the *target* monkey) if we aligned that monkey’s latent signals to those of the source monkey. Fig. 2, left column shows an example of the latent signals obtained by projecting the neural activity of the source and target monkeys into their corresponding neural manifolds (source monkey: J, first session, target monkey: S, first session). The unaligned latent signals of the two monkeys are similar, in that the trajectories corresponding to each of the eight cursor directions are well separated and traverse roughly parallel paths through their respective principal component spaces. However, despite their similar shapes, the two sets of latent signals differ in scale and orientation.

**Fig. 2.**
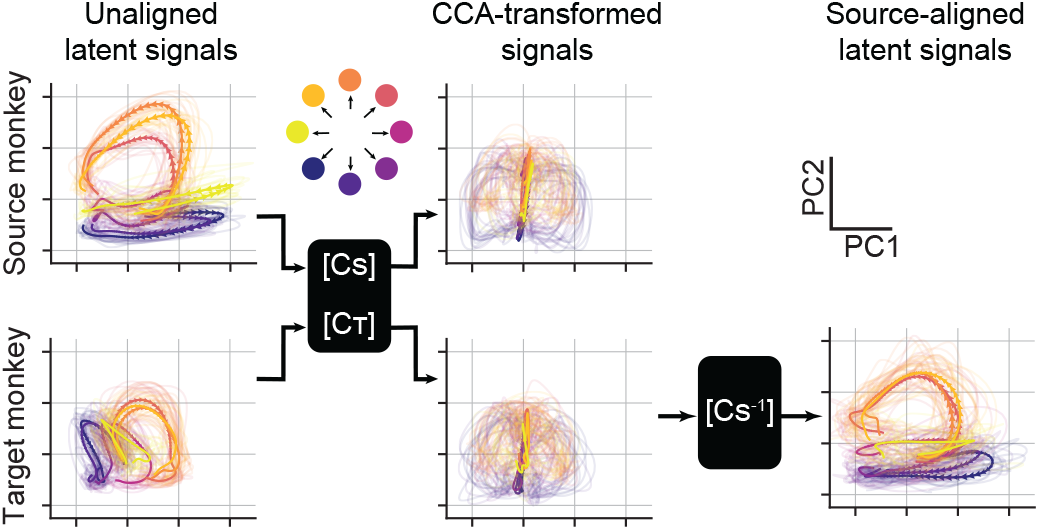
Aligned latent neural signals are very similar across monkeys. Representative latent signals described by the first two principal components for a source (top left) and target (bottom left) monkeys. We used CCA to transform the latent signals such that they were maximally correlated (center). The CCA-transformed signals are very similar to each other, but neither looks like the original source-monkey trajectories. Consequently, we further multiplied the target-monkey latent signals in the CCA-transformed space by the inverse of the source-monkey CCA weights (CS^−1^) to align it to the original source-monkey latent coordinates (bottom right). Data were averaged across all trials for each target direction; single trial trajectories are shown as shaded curves. Arrows indicate the temporal evolution of the trajectories (from

We next used Canonical Correlation Analysis to linearly transform the two sets of low-dimensional signals such that they were maximally correlated (Fig. 2, middle column). Since CCA aims at maximizing the correlation of two sets of signals of equal length, we ensured that the latent signals of the two monkeys were temporally matched by first time-aligning the neural data of each trial with respect to the force onset (see Methods). The transformed sets of latent signals traversed very similar regions of the new (transformed) latent space, but neither looked like the original source-monkey trajectories, as both were transformed by CCA. Consequently, we further processed the transformed target-monkey latent signals by using the inverse of the canonical correlation transformation from the source monkey. This two-step process allowed us to align the target-monkey signals to those of the source monkey (Fig. 2, right column). The aligned trajectories for the two monkeys (Fig. 2, upper left, lower right) are a remarkably close match.

### CCA alignment enables cross-monkey decoding

We used latent signals of the target monkey, once aligned to the signals of the source monkey, as inputs to the decoder trained on source-monkey data in order to predict EMGs (Fig. 3a). If CCA alignment had perfectly matched the motor cortex representations of the target monkey to those of the source monkey, then the decoder should output the same EMG predictions as those of the source monkey. Thus, we compared the target-monkey EMG predictions, as well as well as the original source-monkey predictions, to the actual EMG recordings of the source monkey. In both cases, we determined the EMG decoding accuracy by computing the coefficient of determination (*R*^2^) between the EMG predictions and the actual EMG traces of the source monkey. Choosing the actual EMGs of the source monkey as ground truth is the only option in the case of a paralyzed human user.

**Fig. 3.**
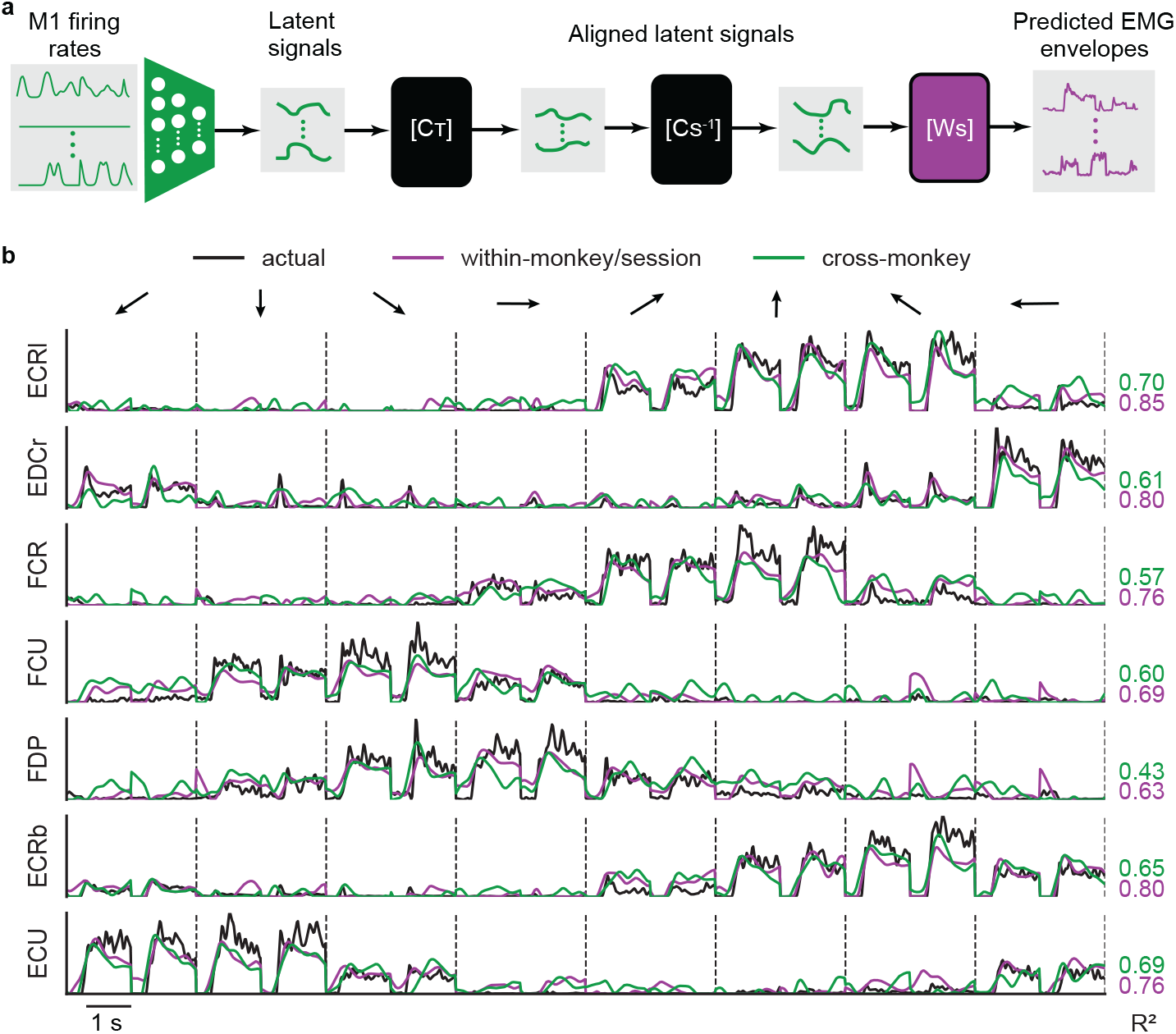
Cross-monkey EMG predictions are similar to within-monkey decoding accuracy after CCA alignment. **a**, Setup for cross-monkey decoding. Neural firing rates from a target monkey performing the isometric wrist task are recorded and projected into their neural manifold to get the corresponding latent signals. CCA is then used to transform these latent signals so that they are maximally correlated with those of the source monkey. The target-monkey latent signals are finally multiplied by the inverse of the source-monkey CCA weights (CS^−1^). The resulting aligned latent signals now provide an input to the source-monkey decoder to obtain predicted EMG activities. **b**, EMG predictions obtained by using the aligned latent signals of the target monkey as inputs to the source-monkey decoder (green lines). These predictions are almost as good as those obtained by using the source-monkey latent signals as input to the decoder trained on data from the source monkey in the same session (purple lines). Actual EMG record-ings of the source monkey (black lines) provide a ground truth for measuring decoding accuracy. The R^2^ for both within-(purple) and cross-(green) monkey decoding are shown on the right for each muscle. Vertical dashed lines separate muscle traces for the eight target directions. Data for a representative pair of mon-keys for two trials is plotted for each target direction.

Fig. 3b shows an example of the within- and cross-monkey EMG decoding accuracy (source monkey: J, first session, target monkey: S, first session). The accuracy of within-monkey EMG decoding ranged between *R*^2^ = 0.63 and 0.85 for individual muscles. The accuracy of cross-monkey decoding was lower, ranging between 0.43 and 0.70 for individual muscles. For some muscles (such as the extensor carpi ulnaris; ECU), the within- and cross-monkey *R*^2^ decoding performances were almost identical (0.76 and 0.69, respectively). The average performance across all muscles, weighted by their individual variances, was 0.76 for the within-monkey predictions and 0.61 for cross-monkey predictions. In this example, the cross-monkey predictions retained 80% of the decoding accuracy of the within-monkey decoders.

We followed the same procedure for all monkey pairs, with similar results (Fig. 4a). The diagonal of this matrix shows the weighted average across all muscles of the within-monkey decoding accuracy within a session; this number sets a strong upper bound on the cross-monkey prediction accuracy and ranged from 0.64 to 0.76. The off-diagonal elements represent either cross-monkey or within-monkey/cross-session accuracy. We used the pipeline shown in Fig. 3a to align the datasets across both. We included the within monkey/cross-session decoding accuracy as a more realistic comparison with the cross-monkey (and implicitly cross-session) analyses. These two cross-session comparisons were not significantly different (within-monkey: 0.54 ± 0.05 (mean ± s.e.); cross-monkey: 0.53 ± 0.02; P = 0.84, Wilcoxon’s rank sum test). Without CCA alignment, cross-monkey predictions resulted in uniformly negative *R*^2^ values (Fig. 4b, right panel). On average, cross-monkey decoding retained 97% of the accuracy of within-monkey (cross-session) decoding, which was 76% even of the within-monkey/within-session decoding accuracy.

**Fig. 4.**
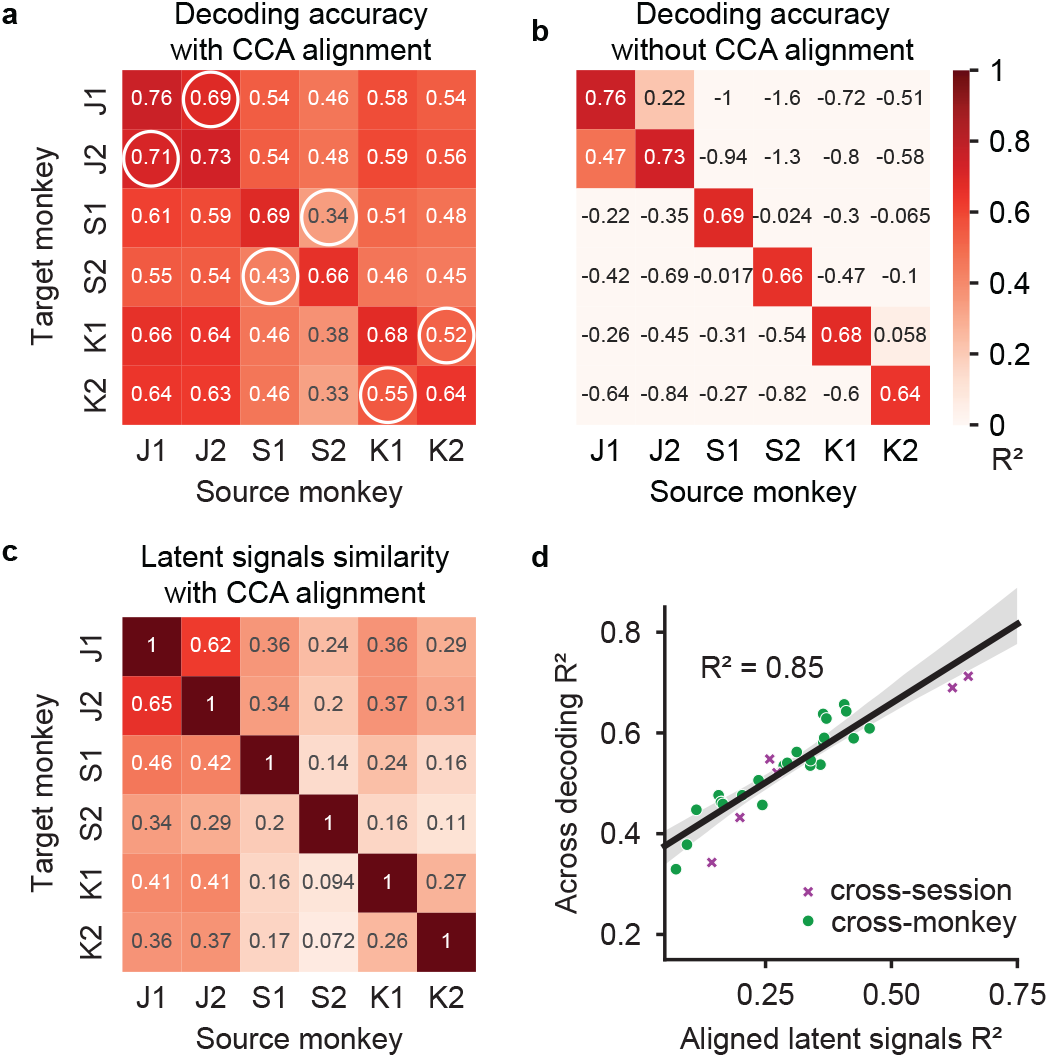
CCA alignment enables accurate cross-monkey decoding for all monkey pairs. **a**, Overall cross-monkey decoding accuracy (R^2^) after CCA alignment for all pairs of monkeys. Circles indicate within-monkey/cross-session predictions. **b**, Cross-monkey R^2^ values obtained if no CCA alignment between the source and the target latent signals is applied. For both **a** and **b**, the x-axis indicates the source-monkey data used to train the decoder, while the y-axis indicates the target-monkey data used to test the decoder. **c**, Similarity of the source (x-axis) and target (y-axis) monkey latent signals after CCA alignment for all monkey pairs. **d**, Element-by-element scatter plot for the matrices in **a** and **c**. Latent signals of monkey pairs with higher similarity after CCA alignment yielded higher decoding accuracy for both cross-monkey and cross-session predictions.

In transferring decoders across monkeys, we could have used the actual EMGs of the target monkey as ground truth, but we chose instead, to treat these predictions as we would need to in the case of a monkey-to-human transfer. In the limit, when the EMG patterns of the two monkeys were identical, this choice would have been irrelevant. EMGs in this relatively simple wrist task were consistent across time for a given monkey (Fig. S2), but less so across monkeys (Fig. S3).

In addition to quantifying the accuracy of the EMG predictions, we also quantified the similarity between latent signals by computing the *R*^2^ between the source- and the target-monkey latent signals before and after CCA alignment. CCA alignment significantly improved the similarity of all the source/target-monkey pairs (*R*^2^ before alignment: −1.02 ± 0.10; after alignment: 0.28 ± 0.02 (mean ± s.e.); P ∼ 0, Wilcoxon’s signed rank test). We hypothesized that the source of the variation in cross-monkey decoding accuracy may be in part due to the varying ability to achieve good alignment with CCA. Fig. 4c shows the similarities between source- and target-monkey latent signals for the same pairs of datasets shown in Fig. 4a. The similarity between this matrix and that in Fig. 4a is evident to the eye and illustrates that success of alignment is predictive of decoder performance. We quantified the relationship by constructing an element-by-element scatter plot of the two matrices (Fig. 4d). Indeed, both the cross-monkey (green symbols) and cross-session (purple symbols) decoding accuracy was well predicted by the success of the CCA alignment, with an *R*^2^ of 0.85 across all datasets. To summarize, when the latent signals could be more closely aligned across either time or monkeys, the decoding accuracy increased proportionately.

### Generalization of cross-monkey decoding

Another important question is how well the decoder performs when the alignment is applied to neural data corresponding to movements beyond those used to derive the alignment transformation. To investigate this issue, we computed the CCA transformation on subsets of movement targets and tested on the remainder. We first tested the ability to interpolate, using data for the four cardinal directions to obtain the CCA transform, then testing this transform on data for the diagonal directions (Fig. 5a). Under these conditions, CCA aligned data generalized well. In contrast, when we required CCA to extrapolate by training it on the four adjacent lower targets and testing it on the upper targets, the alignment failed. While CCA accurately aligns the training data (Fig. 5b, top row), the transformed trajectories towards test targets (Fig. 5b, bottom row) fail to match the corresponding source trajectories.

**Fig. 5.**
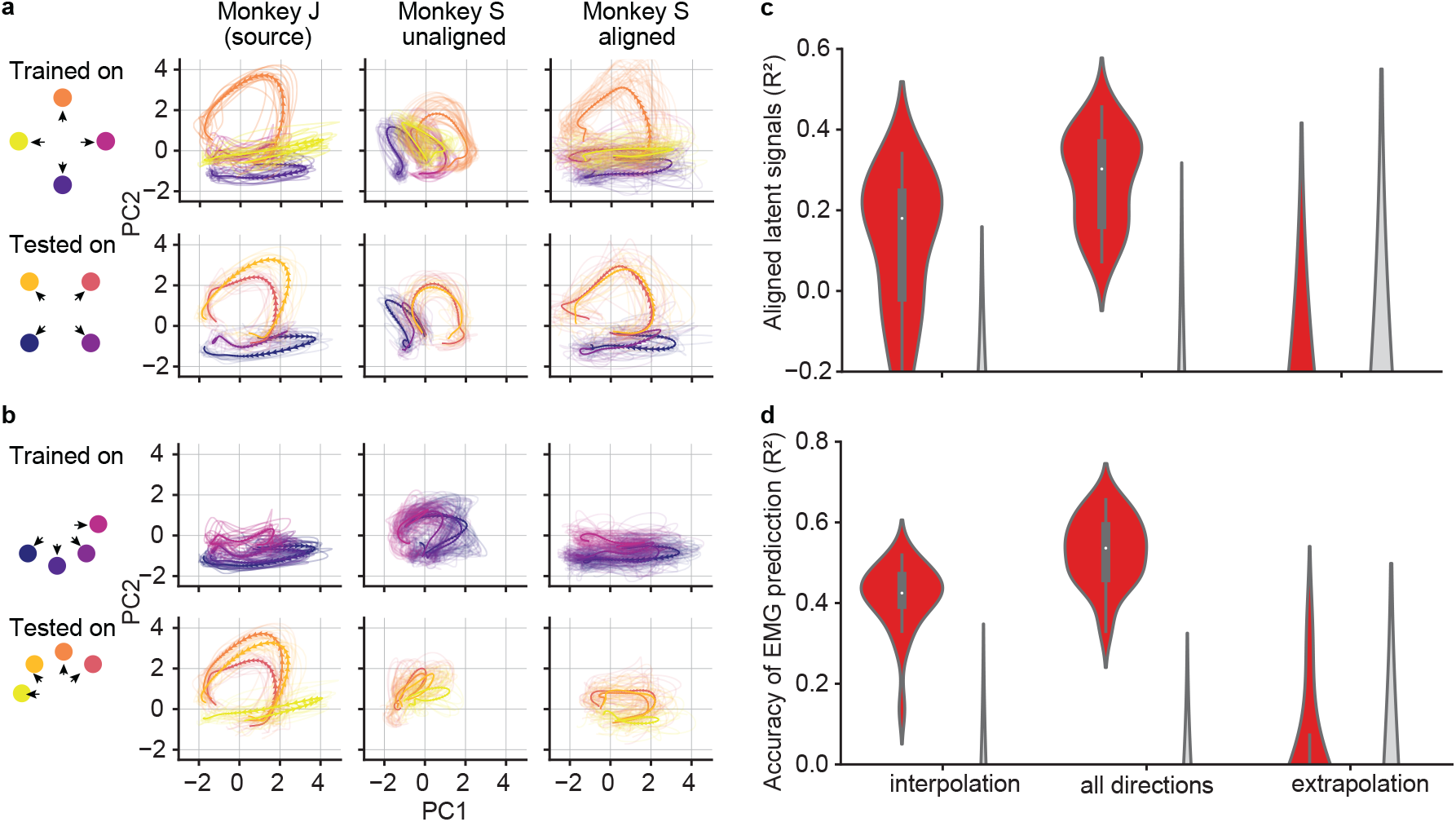
CCA alignment generalizes when interpolating, but not when extrapolating. **a-b**, Representative latent signals described by the first two principal components of a single monkey pair (source monkey: J, first session; target monkey: S, first session) before and after CCA alignment. **a**, Generalizability of the CCA alignment when interpolating (i.e., trained on cardinal directions and tested on diagonal directions); **b**, Generalizability of the CCA alignment when extrapolating (i.e., trained on adjacent lower directions and tested on upper directions). When interpolating, the aligned neural trajectories of the target monkey resembled those of the source monkey when tested on the held-out directions (a, bottom row). When extrapolating, CCA accurately aligned the training data (**b**, top row), but not the trajectories of the held-out directions (b, bottom row). **c**, Violin plot showing the similarity, as measured by R^2^, between the latent signals of all pairs of source and target monkeys before (grey) and after (red) CCA alignment when data from all (center), only cardinal (left), and only lower (right) directions were available. When using all directions and when interpolating, the similarity significantly increased after CCA alignment. When extrapolating, the R^2^ values are still negative even after alignment. **d**, Violin plot for the overall cross-monkey decoding R^2^ for all pairs of monkeys before (grey) and after (red) CCA alignment when testing on all target directions (center), when interpolating (left), and when extrapolating (right). When interpolating, cross-monkey decoding is still possible, albeit with lower accuracy. When extrapolating, the cross-monkey decoding generally failed, as indicated by the negative R^2^ values.

The results across all combinations of datasets were consistent with these examples. The similarity between the latent signals of the two monkeys dramatically increased after the alignment based on the interpolated targets but failing on the extrapolated test set (Fig. 5c). For these three conditions (full training set, interpolated training data, extrapolated training data), the mean ± s.e. values of *R*^2^ between source and test neural signals after CCA alignment were: 0.28 ± 0.02, 0.12 ± 0.03, and −0.99 ± 0.10. The differences between all three groups were significant (P ∼ 0, Wilcoxon’s signed rank test). These results also followed the relation shown in Fig. 4d: cross-monkey EMG decoding *R*^2^ when the alignment was based on all directions was 0.53 ± 0.02. For interpolated and extrapolated alignment, it was 0.41 ± 0.02 and −0.23 ± 0.04, respectively, all significantly different (Fig. 5d, P ∼ 0, Wilcoxon’s signed rank test).

### A monkey decoder can be transferred to a person with tetraplegia

Having demonstrated that a fixed decoder trained on data from one monkey can be transferred to a different monkey through latent-signal alignment, we asked if a similar transfer could be implemented from a nonhuman to a human primate. We thus tried to use a monkey decoder to predict EMG signals from the neural activity of a paralyzed human participant attempting to perform a similar task. Because alignment requires only a relatively small amount of neural data, such an operation might allow humans with paralyzed limbs to use powerful decoders without the need to collect the large amount of data needed for supervised decoder training. For this analysis, we chose to use the first session from monkey J (dataset J1) as the source monkey, as it had the highest within-monkey decoder performance among all datasets.

We performed these experiments with one participant with partial spinal cord injury, who was implanted with 96-channel electrode arrays in both the arm and hand areas of the left primary motor cortex (M1). We recorded multi-unit spiking activity as the participant, who had some remaining ability to produce wrist extension forces, attempted to produce isometric wrist forces in eight radial directions. Cursor control and task design were essentially as in the monkey experiments, with the following exceptions. First, the participant attempted to use their right arm and hand (contralateral to the arrays), thereby reversing flexion and extension muscle activity relative to those of the monkey. Consequently, for CCA computation, we mirrored the target directions for the human data about the vertical axis. Second, cursor movement was not under the control of the participant, but occurred automatically, as in the standard approach to “observation” decoder training ^6^. One second after target appearance, a go cue occurred and the cursor moved to the target in 0.2 s, where it remained for 2.0 s before returning to the center in 0.2 s. We instructed the participant to attempt to produce the forces necessary to control the position of the cursor.

Fig. 6a shows the time course of the neural activity projected onto the first principal component during trials corresponding to two oppositely directed targets. The human data was more variable across trials than that of the monkey, likely due to the lack of actual force and force-related feedback. To minimize the effect of this variability on the alignment process, we computed the correlation across trials paired between the monkey and human data as a function of the time window we used for the CCA alignment. We achieved maximal correlation (expressed by the Pearson’s correlation coefficient averaged across the first five latent dimensions) between the human and monkey latent signals when aligning the human trials 0.76 sec *prior* to the go cue (Fig. 6b) to the time of force onset in the monkey data.

**Fig. 6.**
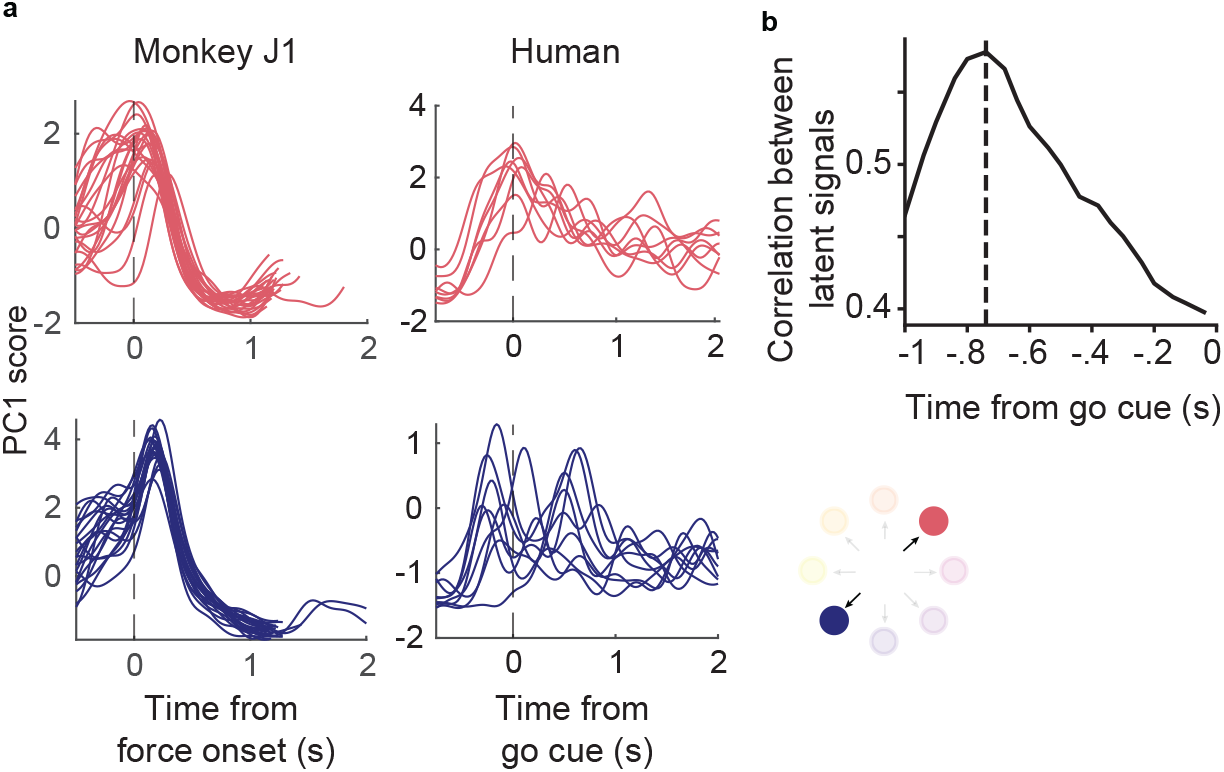
Accuracy of cross-species CCA alignment depends on the latency relative to the go cue used for alignment. **a**, Single-trial M1 data from the source monkey (left, first session from monkey J) and the human participant (right) projected on the first principal component (computed in each case using the entire corresponding dataset) for a pair of oppositely directed targets. The neural responses recorded during the human’s attempted task show increased trial-by-trial variation in their timing when compared to those of the monkey. **b**, Monkey-to-human latent signals correlation, averaged across the first five latent dimensions after CCA alignment as a function of the time alignment. The vertical dashed line indicates the time relative to the go cue that yielded the best correlation.

Despite the greater inter-trial variability, the latent signals of the human data traversed well-separated, stereotypical trajectories for each of the eight targets (Fig. 7a, left panel). CCA alignment allowed us to match the two sets of latent signals fairly closely (Fig. 7a, middle and right panels) improving the similarity to the source monkey from *R*^2^ = −0.56 to *R*^2^ = 0.10. These aligned signals allowed us to make accurate EMG predictions from the human M1 data using a decoder computed from the J1 monkey dataset. The average prediction accuracy for the four major wrist muscles was 0.55 (Fig. 7b, green lines), 89% as accurate as the corresponding cross-monkey predictions, and 72% as accurate even as the average within-monkey/with session decoding (Fig. 7b, purple lines).

**Fig. 7.**
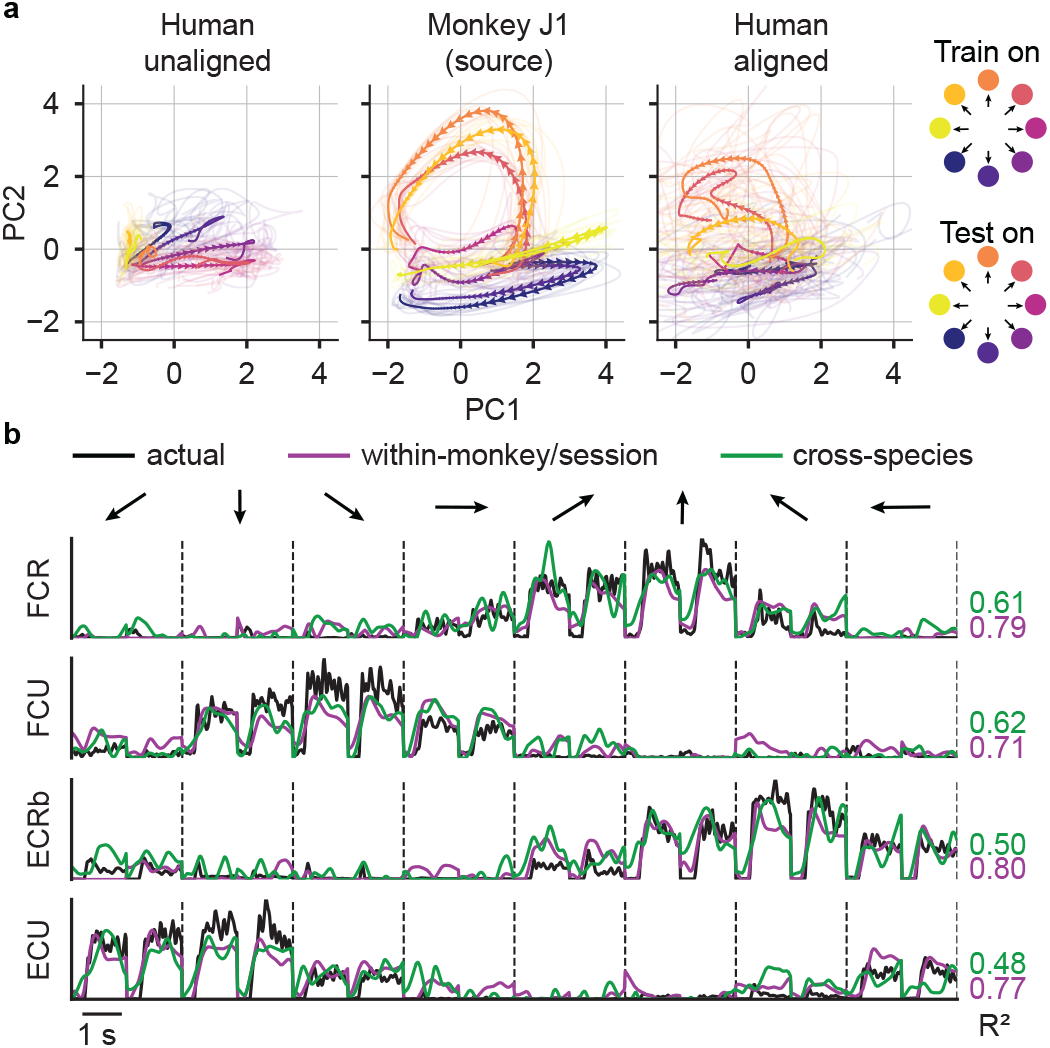
CCA alignment enables the transfer of a monkey decoder to a human with tetraplegia. **a**, Latent M1 trajectories described by the first two principal components of the M1 data (left) recorded as the participant attempted to perform a wrist isometric task, and those for the source monkey (center, first session from monkey J). The human neural recordings show a stereotypical low-dimensional structure for each of the eight target directions, albeit with increased inter-trial variability (shown by the shaded curves). Despite this increased variability, CCA alignment recovered a shape similar to that of the source monkey’s latent signals (right). **b**, EMG predictions for the four major wrist muscles obtained by using the aligned latent signals of the human participant as inputs to the source-monkey decoder (green lines). These predictions are almost as good as EMG predictions obtained by using the source-monkey latent signals and decoder (purple lines). Actual EMG recordings of the source monkey (black) provide a ground truth for measuring decoding accuracy. The R^2^ for both within-(purple) and cross-(green) species decoding are shown on the right for each muscle. Vertical dashed lines separate muscle traces for two trials in each of the eight target directions.

## Discussion

We have demonstrated the transfer of a decoder trained on neural and EMG data from one monkey to another, allowing us to make EMG predictions from the neural activity of the second monkey without needing to compute a new decoder for that monkey. The same methods allowed us to make EMG predictions even from the neural activity recorded during attempted movements from a person with a paralyzed arm. We have used as ground truth for these predictions the EMG signals recorded from the source monkey, as there is no ground truth available for the human participant. We refer to the “task accuracy” of these predictions to reflect this point. The similarity of the human and nonhuman primate musculoskeletal systems means that a muscle-level solution that is effective for one subject is likely to work well for a different individual, even across species. The decoder transfer was possible because the low-dimensional neural representations of motor intent for specific actions were also similar across individuals and species. Our approach relied on “aligning” the latent representations using Canonical Correlation Analysis. We, and others, have used similar methods to align latent representations for a given individual across time. Crucially, this alignment process requires only neural data and in only very limited amounts, making it both practically and theoretically applicable to clinical settings where data collection is difficult and there is no motor output data. Overall, our work suggests that transferring a decoder between monkeys and humans is indeed possible. Our approach can generate EMG predictions that can be used as the control signals for functional electrical stimulation of muscles as a means to restore voluntary arm movement ^5,14,15^ or movement of an anthropomorphic limb actuated with control properties designed to mimic those of the musculoskeletal system.

### Neural representations of motor intent are similar across monkeys

The neural population activity in many brain areas is constrained to a low-dimensional neural manifold. Furthermore, there is increasing evidence that the dynamics of specific patterns of activity within the manifold, the latent signals, underlie the computations required for planning and executing movements ^8–10,19–22^. In the primary motor cortex, recorded latent signals that differ across days can be mathematically transformed (“aligned”) to be more similar to each other. These aligned latent signals maintain a remarkably stable relation to behavior over months and even years ^12^. As a consequence, a fixed decoder that uses these aligned signals as inputs remains accurate across long periods without supervised recalibration ^11,12,23,24^.

Manifolds transcend the analysis of individual neurons, and make the comparisons of population activity across individuals more readily interpretable ^25^. Recently, Safaie et al. used similar alignment techniques to show that latent signals in M1 were preserved across monkeys as they performed a center-out reaching task using a planar manipulandum; latent signals were also preserved in the dorsolateral striatum of mice that grasped and pulled a joystick ^26^. Other recent studies revealed that low-dimensional neural structure within the hippocampus and the sensorimotor cortex is preserved across rats for a variety of behaviors, ranging from locomotion along a linear track inside a maze to unconstrained movement in an arena ^27–30^.

It is important to note that as our monkeys adopted differing muscle activation strategies during the task (possible because of the muscle-level redundancy of the limb), the neural representations of the task also became more dissimilar (Fig. S5d). The magnitude of the relationship between M1 and EMG similarity was reasonably high (*R*^2^ = 0.4), though not nearly as high as that between the similarity of the M1 latent signals and the corresponding decoding accuracy (*R*^2^ = 0.85, Fig. 4d). If M1 encoded a task specific motor intent signal that was equivalent across monkeys but implemented via differing muscle synergies, the relationship shown in Fig. S5d would have been flat. However, the fact that M1 alignment is a much better predictor of EMG-prediction accuracy (Fig. 4d) than it is of EMG-similarity suggests that at least a component of neural code was preserved despite different task execution strategies. This observation may be consistent with the degree of task independence seen in M1 in several earlier studies ^31,32^.

### Using the preserved motor intent signal to transfer monkey decoder to a human

Not only was the neural representation of the isometric wrist task similar across monkeys, but the similarity extended even to a paralyzed human attempting to perform the same task. Remarkably, the EMG predictions obtained with the human M1 data still resembled the actual EMGs of the source monkey, even though the decoding accuracy was not as high as when using M1 data from a target monkey. The neural trajectories derived from the human M1 data had greater inter-trial variability compared to those of the monkeys (Fig. 6a), but they were still well separated for each of the eight target directions (Fig. 7a). The greater variability in the human M1 recordings was undoubtedly due at least in part to the lack of any actual force or force-related feedback to the participant during the attempted movements. The variability may also have been due to changes in M1 in the 10 years since the participant’s injury.

The lack of actual force output also posed a challenge for transferring a decoder from monkey to humans as there was no clear onset time for each attempted movement. In the monkey datasets we time-aligned each trial to the onset of force. With the human data, we had to rely on a consistent reaction time with respect to the go cue. For at least some targets, the participant appeared to anticipate the go cue (Fig. 6a). As a result, the accuracy of the CCA alignment peaked for time-alignment *prior to* the go cue (Fig. 6b).

Unlike the single array in the hand area of monkeys M1, the human participant had separate arrays implanted in both the hand and the arm areas. As expected, *R*^2^ was roughly 30% higher when we decoded EMG from the hand array compared to the arm array (Fig. S8b). The neural trajectories for the arm area were also reasonably well separated across targets, but their average firing rates were lower than those of the hand area array, as indicated by the smaller magnitude of the activity projected onto the first two PCs (Fig. S8a). The combined use of arm and hand neural recordings led to slightly higher performance (Fig. S8b). The ability to predict wrist and hand EMGs from the arm area array could be due to co-contraction of arm and hand muscles as the participant attempted to produce wrist force, or it might be the result of hand-related inputs to the arm-area neurons. A similar observation about mixed representations has been made in a study with different human participants, for whom it was possible to predict attempted movement of a broad range of body parts from recordings made in the hand area of M1 ^33^.

### Clinical applicability of monkey-to-human transfer of a biomimetic decoder

While existing iBCIs have allowed paralyzed individuals to gain closed-loop control of a computer cursor with only a few minutes of practice ^34^, higher-dimensional control is more difficult for the user to learn ^2,4^. Furthermore, existing iBCIs allow no direct control of applied grasp forces. Attempts to combine position and joint torque control for 2D planar reaching have met with limited success ^35,36^. These limitations are despite the well documented evidence of force and other kinetic information in the primary motor cortex ^37–46^.

The mammalian neuromuscular system controls the motion of the arm and digits, the stiffness of joints, and exerted forces, all through the modulation of muscle activity. One approach to a more biomimetic form of robotic limb control allowing more intuitive control of many degrees of freedom would be to feed real-time predictions of muscle activity to a musculoskeletal model that would compute muscle forces and the evolution of limb state; these signals could be used to control the motion and impedance of an anthropomorphic prosthetic arm ^47–49^.

Alternatively, the predicted EMG signals could be used to control muscle force directly through the electrical stimulation of the muscles or peripheral nerves, a technique known as Functional Electrical Stimulation (FES). Our group previously designed an iBCI-controlled FES system that enabled monkeys with temporary paralysis of the hand muscles induced by a peripheral nerve block to perform hand movements: decoded EMGs were used to modulate stimulation of five electrodes implanted in different compartments of three hand flexor muscles ^5^.

Two other groups have developed brain-controlled FES systems in which human participants with cervical spinal cord injury were able to control simple elbow, wrist, or hand movements. One approach used six parallel decoders, each predicting one of six wrist or finger movements. The decoder with the highest output triggered a muscle stimulation pattern consisting of at most, three intensity levels, designed to approximate the decoded movement ^15^. In another approach, real-time velocities of the elbow, wrist, hand, and shoulder were decoded from M1, and controlled in a feedback system using a combination of muscle and nerve stimulation ^14^.

While promising as additional proofs of concept, these approaches achieved only limited control of a small number of dimensions. The former, in particular, seems unlikely to be able to scale to more complex movements. In our study, we aimed to achieve the means to provide more intuitive control of more degrees of freedom (including nonkinematic aspects of movement) by directly inferring muscle activity from M1.

### Limitations and future directions

CCA is fundamentally limited in that it is a linear method for which the two datasets need to correspond to the same task condition and be temporally aligned. While our task allowed us to time-align the monkey data, the same was not possible for the human data, potentially limiting the effectiveness of the alignment. A different linear approach has been used to directly align the coordinate axes of manifolds across sessions that does not rely on temporal alignment of the data ^11^. This method, however, relies on the existence of at least a subset of stably recorded neurons that anchor the realignment, making it unlikely to be useful to align recordings from different individuals for which there can be no neurons in common.

Other techniques have been used to align the statistics of two clouds of points, independently of the time course of the associated signals ^23,24,50^. These unsupervised methods have been tested on neural recordings corresponding to structured, trial-based behaviors, although they have been successful even without the use of any information about the behavior itself. To our knowledge, they have not been applied to the highly varied and unstructured movements typical of the activities of daily living. These and related methods should be further investigated under more realistic conditions.

As noted above, the redundancy of the hand musculature allows most actions to be produced with a variety of different patterns of muscle activity. If it were the case that the human participant intended to use a different pattern of EMG from that of the source monkey, CCA would nonetheless attempt to make the corresponding neural trajectories as similar as possible. To the extent that the biomechanics of the human hand resembles that of the monkey, the resultant EMG predictions should be biomechanically appropriate. However, it could pose an interesting motor-adaptation problem as the user learns to interact with the decoder.

A further question is the extent to which a decoder trained in a limited setting might generalize to a broader range of motor actions. We have at least preliminary evidence that simple linear decoders do not generalize well ^51,52^. Here is where the real strength of the decoder transfer may lie. While alignment required very little data (see Methods), a complex, nonlinear decoder capable of broad generalization is likely to require more data than is feasible to obtain directly from a person with a paralyzed limb. Sussillo et al., showed that a decoder trained with data recorded from a monkey over sessions spanning as many as 50 days significantly improved decoding accuracy and iBCI performance ^53^. While this would likely not be feasible with a human user, training a decoder with a large amount of monkey data prior to augmentation with a small amount of subject-specific data may indeed be a powerful approach, which we have not yet begun to explore.

## Methods

### Monkey task and recordings

Neural and muscle activity data were collected from three adult male rhesus macaque monkeys. All surgical and experimental procedures were approved by the Institutional Animal Care and Use Committee (IACUC) of Northwestern University.

The monkeys were seated in a primate chair that faced a computer monitor, with their left hand secured within a small box instrumented with a 6 degree of freedom load cell (JR3 Inc., CA) to measure the forces exerted (Fig. 1a). The load cell measurement axes were aligned with those of the wrist so that flexion/extension forces moved a cursor on the monitor right/left, while radial/ulnar deviation forces moved it up/down. The monkeys were required to move the cursor from a central target towards one of eight peripheral targets uniformly distributed on a circle. A trial started when the monkeys held the cursor in the central target for a random time between 0.2 s and 1.0 s. Then, one of the eight peripheral targets was selected randomly and presented together with an auditory go cue. Monkeys had to reach the outer target within 2.0 s and maintain that force for 0.8 s to receive a liquid reward. In this study, we used data within a window starting 0.5 s before onset of cursor movement and ending 1.0 s after movement onset.

Each monkey was implanted with a 96-channel Utah electrode array (Blackrock Neurotech, Inc.) in the hand area of the right M1, contralateral to the hand used for the task, using standard surgical procedures. In a separate procedure, monkeys were also implanted with intramuscular leads in the forearm and hand muscles of the left arm. The location of each electrode was verified during the surgery by observing the muscle contraction evoked when it was stimulated. Five muscles were implanted in all three monkeys: three major wrist muscles: extensor carpi radialis longus (ECRl), flexor carpi radialis (FCR), and flexor carpi ulnaris (FCU), and two extrinsic finger muscles: extensor digitorum communis radialis (EDCr) and flexor digitorum profundus (FDP). Beside these muscles that were in common across monkeys, we implanted additional muscles in each monkey. These included extensor carpi radialis brevis (ECRb) and extensor carpi ulnaris (ECU) in monkey J, flexor digitorum indicis (FDI) and opponens digiti minimi (MD) and FDP (2 pairs) in monkey S, and ECU, extensor digitorum communis (EDC), flexor digitorum superficialis (FDS), pronator teres (PT), and supinator (SUP) in monkey K.

We recorded M1 activity using a Cerebus system (Blackrock Neurotech, Inc.). The signals on each channel were sampled at 30 kHz, digitally bandpass filtered (250 ∼ 5000 Hz) and converted to spike times based on threshold crossings. The threshold on each channel was set with respect to its root-mean square (RMS) amplitude (monkey J: −5.5 x RMS; monkey S: −6.25 x RMS; monkey K: −6.0 x RMS). To extract the smoothed firing rate used in the analysis, we applied a Gaussian kernel (standard deviation of 100 ms) to the spike counts in 20 ms, non-overlapping bins for each channel.

The recorded EMG signals were amplified, bandpass filtered (4-pole, 50 ∼ 500 Hz), and sampled at 2 kHz. To extract the envelopes used during the analysis, we subsequently rectified and lowpass filtered (4-pole Butterworth, 10 Hz) each EMG channel digitally and subsampled it to 50 Hz to correspond to the bin size of the M1 signals. EMGs were clipped to avoid data points larger than the mean plus 6 times the standard deviation of each channel. We removed the baseline of each EMG channel by subtracting the 2^nd^ percentile of its amplitude and then normalized its activity to its 90^th^ percentile.

### Human participant tasks and recordings

The participant, male, 28 years old at the time of the implant, was part of a multi-site clinical trial (NCT01894802) and provided informed consent prior to the experimental procedure. He presented with a C5 motor/C6 sensory ASIA B spinal cord injury that occurred 10 years prior. He had no spared control of the intrinsic or extrinsic muscles of the right hand but had limited control of wrist flexion and extension. Proximal limb control at the shoulder was intact, as was elbow flexion. However, he had no voluntary control of elbow extension.

We secured a rigid wrist brace to the participant’s right wrist and hand, oriented in a neutral posture on his lap. On each trial, the participant attempted to produce isometric wrist forces in response to the movement of a cursor to eight different radial targets displayed on a screen in front of him. Upward movement corresponded to radial deviation, rightward movements to wrist extension, downward movements to ulnar deviation, and leftward movements to wrist flexion. Diagonal targets corresponded to the appropriate combinations of these gestures. Each trial began with the presentation of the upcoming target. One second after target appearance, a go cue occurred, followed by movement of the cursor to the target (Move: 0.2 s), static hold at the target (Hold: 2.0 s), and return to center (Return: 0.2 s). We instructed the participant to produce step-like force profiles in response to the movement of the cursor. To match the time scale of the monkey data, here we only used data from a 1.5 s window, starting 0.76 s before and ending 0.74 s after the go cue (see Fig. 6b).

The participant was implanted with four NeuroPort microelectrode arrays (Blackrock Neurotech, Inc.) in the left hemisphere. Two 96-channel arrays were implanted in the hand and arm area of M1, while the other two were implanted in somatosensory cortex. In this study, we only used the arrays placed in M1. The signals on each channel were recorded at 30 kHz and high-pass filtered at 750 Hz. Whenever the signal crossed a threshold (−4.25 x RMS), a spiking event was recorded. We used multiunit threshold crossing and considered spike counts in 20 ms non-overlapping bins. We applied a Gaussian kernel (standard deviation of 100 ms) to obtain the smoothed firing rate used in the analysis. For all data analyses, we mirrored the target labels for the human data as his cortical implant is in the opposite hemisphere as that of the monkeys, who used their left hands to perform the task.

### Fixed iBCI decoder

We computed EMG predictions using a Weiner cascade decoder on inputs provided by latent signals within a low-dimensional manifold of M1. The Wiener cascade ^18^ consists of a linear filter followed by a static nonlinearity. The Wiener filter implemented linear regression to predict the EMG at the current time bin from the neural recordings stretching to five time bins (100 ms) into the past. The output of the linear filter was then rectified.

To find the latent signals, we performed dimensionality reduction by applying PCA to the M1 firing rates. We estimated the linear dimensionality of the M1 data of each monkey using Parallel Analysis (PA) and set the dimensionality of the latent space to 13, the largest estimate across all monkeys and sessions. PA is a noise-robust linear dimensionality estimator method that finds the number of significant eigenvalues through comparisons to the null distribution for each eigenvalue obtained from repeated, independent shuffling of the data over time within each channel. We have used PA to estimate the dimensionality of both simulated and actual neural data (see ^16^).

We trained the Wiener cascade filter using data from a given monkey (the *source monkey*) with a 4-fold cross validation. For each monkey, we used a total of 16 trials for each of the eight target directions in the cross-validation, yielding a total of 96 trials as the training set and 32 trials as the test set for each fold. We selected the decoder for the fold that yielded the best *R*^2^ on the test set as the fixed iBCI decoder for that monkey. Since we recorded two sessions for each monkey, we trained six source-monkey decoders.

### CCA alignment of low-dimensional neural signals

In this study, we hypothesized that transferring a neural decoder across users would be possible if the corresponding low-dimensional neural signals of the users were similar. After training the fixed decoder on the latent signals of the source monkey, our goal was to apply it to latent signals derived from the neural recordings of a target monkey or a human with a paralyzed arm. We used PCA to find latent neural spaces for both monkeys (and the human) and defined that of the source monkey as ***S*** and that of the target monkey/human as ***T***. Both are *M x p* matrices, where *M* is the number of samples and *p* the latent dimensionality (*p=13*). We used Canonical Correlation Analysis (CCA) to make ***T*** as similar as possible to ***S***. CCA aims at (linearly) transforming both ***S*** and ***T*** such that the correlation between *C*_*S*_(***S***) and *C*_*T*_(***T***) is maximal, where *C*_*S*_ and *C*_*T*_ are the matrices that implement the linear transformations of the latent signals (Fig. 2). In other words, CCA finds alternative coordinate axes for ***S*** and ***T*** such that the corresponding latent signals are maximally correlated. For latent variables obtained by PCA, the transformations *C*_*S*_ and *C*_*T*_ consist of a rotation followed by a nonisotropic rescaling, which preserves their orthogonality.

Since the EMG decoder is trained on the latent signals in the PC embedding space, ***S***, of the source monkey, we needed to represent ***T*** in that space, not in the transformed space *C*_*T*_(***T***). To achieve this, we further multiplied *C*_*T*_(***T***) by the inverse 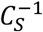 of the source-monkey CCA transformation in order to bring the target-monkey/human latent space coordinates into the original PC embedding space of the source monkey. We refer to the resulting latent signals 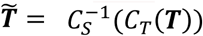 as the aligned target latent signals. 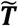 can now be used as input to the fixed source-monkey decoder (Fig. 3a).

We obtained the CCA transformation matrices using k-fold cross validation. Since CCA requires a one-to-one correspondence between data points in ***S*** and ***T***, each training fold consisted of the time-aligned, concatenated latent signals of single trials. For the cross-monkey analysis we used 4-folds, each consisting of 4-concatenated trials for each direction, for a total of 32 trials. Because the amount of data available for the human participant was limited, the monkey-to-human analysis was based on only three folds, each consisting of three trials for each of the eight targets.

### Cross-subject decoding

After training a *source-monkey* decoder with the latent signals of one of the six datasets (three monkeys, two sessions per monkey), we computed the CCA alignment between the latent signals of the source monkey and those of the *target monkey* for each of the remaining five datasets, as described above. Note that of these five datasets, four were from a monkey that was not used to train the decoder, while one was from the same monkey used to train the decoder but recorded during a different session. We tested each of the six source-monkey decoders using the held-out aligned latent signals of a target monkey.

For the monkey-to-human decoder transfer, we performed CCA between the latent signals of the single source monkey with the highest within-monkey decoder performance (dataset J1: monkey J, first session) and those of the paralyzed human participant. We then tested the J1 decoder using the aligned human latent signals.

We compared the resulting EMG predictions with the actual EMG recordings of the source monkey (i.e., the EMG data used to train the fixed decoder). Ideally, the accuracy of the cross-monkey and cross-species decoder as measured by *R*^2^ would approach the performance of the corresponding within-monkey decoder. As the decoded EMGs are multi-dimensional, we computed an *R*^2^ for each EMG and then reported the average across all muscles, weighted by the variances of each muscle. We used the “r2_score” function of the scikit-learn python package with “variance weighted” for the “multioutput” parameter ^54^.

### Generalization of CCA alignment

For the generalization analysis, we computed CCA on a subset of target directions (i.e., “train partition”) and used the remainder target directions (i.e., “test partition”) to assess the quality of alignment and prediction. In the interpolation condition, the train partition considered targets from the cardinal directions, while the test partition considered those from the diagonal directions. In the extrapolation condition, the train partition considered four adjacent lower targets, and the test partition the remaining adjacent upper ones (Fig. 5b). In both conditions, the decoder was computed using source-monkey data for all targets.

The train and test partition for the source and the target monkeys’ unaligned latent signals were obtained differently. For the source monkey, we computed PCA on M1 data from all directions and then split the resulting latent projections into the train and test partitions according to the related target directions. For the target monkey, we computed PCA only on M1 data from a subset of directions (i.e., the directions of the train partition). Then, we projected the M1 data of the other subset of directions (i.e., directions of the test partition) using the PCA weights of the train partition.

## Acknowledgments

We thank Eric J. Perrault for valuable discussions. We thank current and former members of the Miller Limb Lab, including Stephanie Naufel and Emily Oby, for their contributions to data collection. The work was supported in part by grants to L.E.M. (R01 NS053603, R01 NS074044).

## Author Contributions

F.R., E.A., and L.E.M. designed research; X.M and K.L.B. performed experiments; F.R. and E.A. analyzed data; F.R. and E.A. drafted manuscript, X.M., K.L.B., S.A.S., A.K., and L.E.M. edited and revised manuscript; and S.A.S., A.K., and L.E.M. provided supervision.

## REFERENCES

1. Carmena, J. M. et al. Learning to Control a Brain–Machine Interface for Reaching and Grasping by Primates. PLOS Biology 1, e42 (2003).

2. Collinger, J. L. et al. High-performance neuroprosthetic control by an individual with tetraplegia. The Lancet 381, 557–564 (2013).

3. Dekleva, B. M., Weiss, J. M., Boninger, M. L. & Collinger, J. L. Generalizable cursor click decoding using grasp-related neural transients. J. Neural Eng. 18, 0460e9 (2021).

4. Wodlinger, B. et al. Ten-dimensional anthropomorphic arm control in a human brainmachine interface: difficulties, solutions, and limitations. J. Neural Eng. 12, 016011 (2014).

5. Ethier, C., Oby, E. R., Bauman, M. J. & Miller, L. E. Restoration of grasp following paralysis through brain-controlled stimulation of muscles. Nature 485, 368–371 (2012).

6. Willett, F. R. et al. Principled BCI Decoder Design and Parameter Selection Using a Feedback Control Model. Sci Rep 9, 8881 (2019).

7. Churchland, M. M. et al. Neural population dynamics during reaching. Nature 487, 51–56 (2012).

8. Cunningham, J. P. & Yu, B. M. Dimensionality reduction for large-scale neural recordings. Nat Neurosci 17, 1500–1509 (2014).

9. Gao, P. & Ganguli, S. On simplicity and complexity in the brave new world of large-scale neuroscience. Current Opinion in Neurobiology 32, 148–155 (2015).

10. Gallego, J. A., Perich, M. G., Miller, L. E. & Solla, S. A. Neural Manifolds for the Control of Movement. Neuron 94, 978–984 (2017).

11. Degenhart, A. D. et al. Stabilization of a brain–computer interface via the alignment of lowdimensional spaces of neural activity. Nat Biomed Eng 4, 672–685 (2020).

12. Gallego, J. A., Perich, M. G., Chowdhury, R. H., Solla, S. A. & Miller, L. E. Long-term stability of cortical population dynamics underlying consistent behavior. Nat Neurosci 23, 260–270 (2020).

13. Bach, F. R. & Jordan, M. I. Kernel Independent Component Analysis. Journal of Machine Learning Research 3, 1–48 (2002).

14. Ajiboye, A. B. et al. Restoration of reaching and grasping movements through brain-controlled muscle stimulation in a person with tetraplegia: a proof-of-concept demonstration. Lancet 389, 1821–1830 (2017).

15. Bouton, C. E. et al. Restoring cortical control of functional movement in a human with quadriplegia. Nature 533, 247–250 (2016).

16. Altan, E., Solla, S. A., Miller, L. E. & Perreault, E. J. Estimating the dimensionality of the manifold underlying multi-electrode neural recordings. PLoS Comput Biol 17, e1008591 (2021).

17. Buja, A. & Eyuboglu, N. Remarks on Parallel Analysis. Multivariate Behav Res 27, 509–540 (1992).

18. Hunter, I. W. & Korenberg, M. J. The identification of nonlinear biological systems: Wiener and Hammerstein cascade models. Biol. Cybern. 55, 135–144 (1986).

19. Elsayed, G. F. & Cunningham, J. P. Structure in neural population recordings: an expected byproduct of simpler phenomena? Nat Neurosci 20, 1310–1318 (2017).

20. Mante, V., Sussillo, D., Shenoy, K. V. & Newsome, W. T. Context-dependent computation by recurrent dynamics in prefrontal cortex. Nature 503, 78–84 (2013).

21. Mazor, O. & Laurent, G. Transient dynamics versus fixed points in odor representations by locust antennal lobe projection neurons. Neuron 48, 661–673 (2005).

22. Williamson, R. C., Doiron, B., Smith, M. A. & Yu, B. M. Bridging large-scale neuronal recordings and large-scale network models using dimensionality reduction. Curr Opin Neurobiol 55, 40–47 (2019).

23. Karpowicz, B. M. et al. Stabilizing brain-computer interfaces through alignment of latent dynamics. http://biorxiv.org/lookup/doi/10.1101/2022.04.06.487388 (2022) doi:10.1101/2022.04.06.487388.

24. Ma, X., Rizzoglio, F., Perreault, E. J., Miller, L. E. & Kennedy, A. Using adversarial networks to extend brain computer interface decoding accuracy over time. http://biorxiv.org/lookup/doi/10.1101/2022.08.26.504777 (2022) doi:10.1101/2022.08.26.504777.

25. Dabagia, M., Kording, K. P. & Dyer, E. L. Comparing high-dimensional neural recordings by aligning their low-dimensional latent representations. Preprint at http://arxiv.org/abs/2205.08413 (2022).

26. Safaie, M. et al. Preserved neural population dynamics across animals performing similar behaviour. 2022.09.26.509498 Preprint at https://doi.org/10.1101/2022.09.26.509498 (2022).

27. Chen, H.-T., Manning, J. R. & van der Meer, M. A. A. Between-subject prediction reveals a shared representational geometry in the rodent hippocampus. Current Biology 31, 4293-4304.e5 (2021).

28. Melbaum, S. et al. Conserved structures of neural activity in sensorimotor cortex of freely moving rats allow cross-subject decoding. 2021.03.04.433869 Preprint at https://doi.org/10.1101/2021.03.04.433869 (2022).

29. Nieh, E. H. et al. Geometry of abstract learned knowledge in the hippocampus. Nature 595, 80–84 (2021).

30. Rubin, A. et al. Revealing neural correlates of behavior without behavioral measurements. Nat Commun 10, 4745 (2019).

31. Gallego, J. A. et al. Cortical population activity within a preserved neural manifold underlies multiple motor behaviors. Nat Commun 9, 4233 (2018).

32. Kobak, D. et al. Demixed principal component analysis of neural population data. eLife 5, e10989 (2016).

33. Willett, F. R., Avansino, D. T., Hochberg, L. R., Henderson, J. M. & Shenoy, K. V. Highperformance brain-to-text communication via handwriting. Nature 593, 249–254 (2021).

34. Brandman, D. M. et al. Rapid calibration of an intracortical brain–computer interface for people with tetraplegia. J. Neural Eng. 15, 026007 (2018).

35. Chhatbar, P. Y. & Francis, J. T. Towards a Naturalistic Brain-Machine Interface: Hybrid Torque and Position Control Allows Generalization to Novel Dynamics. PLOS ONE 8, e52286 (2013).

36. Fagg, A. H., Ojakangas, G. W., Miller, L. E. & Hatsopoulos, N. G. Kinetic trajectory decoding using motor cortical ensembles. IEEE Trans Neural Syst Rehabil Eng 17, 487–496 (2009).

37. Cheney, P. D. & Fetz, E. E. Functional classes of primate corticomotoneuronal cells and their relation to active force. Journal of Neurophysiology 44, 773–791 (1980).

38. Evarts, E. V. Relation of pyramidal tract activity to force exerted during voluntary movement. J Neurophysiol 31, 14–27 (1968).

39. Hepp-Reymond, M.-C., Hüsler, E. J., Maier, M. A. & Qi, H.-X. Force-related neuronal activity in two regions of the primate ventral premotor cortex. Can. J. Physiol. Pharmacol. 72, 571– 579 (1994).

40. Holdefer, R. N. & Miller, L. E. Primary motor cortical neurons encode functional muscle synergies. Exp Brain Res 146, 233–243 (2002).

41. Kalaska, J. F. & Hyde, M. L. Area 4 and area 5: differences between the load direction-dependent discharge variability of cells during active postural fixation. Exp Brain Res 59, 197–202 (1985).

42. Lemon, R. N., Johansson, R. S. & Westling, G. Corticospinal control during reach, grasp, and precision lift in man. J. Neurosci. 15, 6145–6156 (1995).

43. Maier, M. A., Bennett, K. M., Hepp-Reymond, M. C. & Lemon, R. N. Contribution of the monkey corticomotoneuronal system to the control of force in precision grip. Journal of Neurophysiology 69, 772–785 (1993).

44. Morrow, M. M., Jordan, L. R. & Miller, L. E. Direct Comparison of the Task-Dependent Discharge of M1 in Hand Space and Muscle Space. Journal of Neurophysiology 97, 1786– 1798 (2007).

45. Oby, E. R., Ethier, C. & Miller, L. E. Movement representation in the primary motor cortex and its contribution to generalizable EMG predictions. Journal of Neurophysiology 109, 666–678 (2013).

46. Sergio, L. E. & Kalaska, J. F. Systematic Changes in Motor Cortex Cell Activity With Arm Posture During Directional Isometric Force Generation. Journal of Neurophysiology 89, 212–228 (2003).

47. Blana, D., Chadwick, E. K., van den Bogert, A. J. & Murray, W. M. Real-time simulation of hand motion for prosthesis control. Computer Methods in Biomechanics and Biomedical Engineering 20, 540–549 (2017).

48. Blana, D. et al. Model-Based Control of Individual Finger Movements for Prosthetic Hand Function. IEEE Transactions on Neural Systems and Rehabilitation Engineering 28, 612– 620 (2020).

49. McFarland, D. C. et al. A Musculoskeletal Model of the Hand and Wrist Capable of Simulating Functional Tasks. 2021.12.28.474357 Preprint at https://doi.org/10.1101/2021.12.28.474357 (2021).

50. Farshchian, A. et al. Adversarial Domain Adaptation for Stable Brain-Machine Interfaces. Preprint at http://arxiv.org/abs/1810.00045 (2019).

51. Ma, X., Bodkin, K. L. & Miller, L. E. Population Activity in Motor Cortex is Influenced by the Contexts of the Motor Behavior. in 2021 10th International IEEE/EMBS Conference on Neural Engineering (NER) 1152–1155 (2021). doi:10.1109/NER49283.2021.9441430.

52. Naufel, S., Glaser, J. I., Kording, K. P., Perreault, E. J. & Miller, L. E. A muscle-activity-dependent gain between motor cortex and EMG. J Neurophysiol 121, 61–73 (2019).

53. Sussillo, D., Stavisky, S. D., Kao, J. C., Ryu, S. I. & Shenoy, K. V. Making brain–machine interfaces robust to future neural variability. Nat Commun 7, 13749 (2016).

54. Pedregosa, F. et al. Scikit-learn: Machine Learning in Python. J. Mach. Learn. Res. 12, 2825–2830 (2011).

