## Supplementary Information for "Monkey-to-human transfer of brain-computer interface decoders"

**This PDF file includes:**

Supplementary text
Supplementary figures S1 to S8

### Supplementary results

#### Monkeys employ slightly different muscle strategies for the same task.

Although the task was the same, there was no guarantee that the three monkeys used similar muscle-level strategies. Moreover, the performance when controlling the cursor differed slightly across monkeys (Fig. S1), potentially resulting in further differences in their muscle strategies. To the extent the recorded EMG activity differs across monkeys, the EMG prediction accuracy of the target monkey will suffer.

We computed the correlation patterns for all recorded muscles for both sessions (Fig. S2a), and observed consistent patterns over time for all monkeys (Fig. S2b). We quantified the difference in muscle activations across two different recording sessions for each monkey by computing their tuning curves and preferred directions over the 1.5 seconds trial-window. Each monkey adopted consistent muscle activation patterns over time, as the change in the PDs ($\Delta PD$, reported as absolute value) recorded across the two sessions was small for all muscles (Table S1, Fig. S2b bottom row).

Then, we quantified the similarity of muscle-level strategies across monkeys. First, we compared the trial-averaged EMG activity of muscles that were recorded in common across all three monkeys (Fig. S3a). While most muscle activation patterns were fairly consistent across monkeys, we also observed some individualized strategies. FCR and EDC differed the most across monkeys (Fig. S3b, Table S1). The activation of flexor carpi radialis for Monkey K (FCR, orange lines in Fig. S3a) was higher than the other two monkeys for the right and down targets, co-contracting it along with flexor carpi ulnaris (FCU, also orange lines). Monkey J activated extensor digitorum communis (EDC, purple lines) with a brief, phasic burst around force onset for most target directions, in contrast with the more tonic activation employed by monkeys S (green lines) and K (orange lines). We also compared the correlations of such muscles across monkeys (Fig. S3c) and observed a similar result, with EDC having substantially to moderately low correlation between monkey J and S ($r = 0.06$) and Monkey J and K ($r = 0.41$), respectively. Similarly, FCR was only moderately correlated between Monkey S and K ($r = 0.39$). All other muscles are highly correlated ($r > 0.6$) across monkeys.

#### Generalization of cross-species decoding.

As for the cross-monkey analysis, we also tested the robustness of the CCA alignment approach for the cross-species decoding when generalizing to task conditions other than those used for training CCA. When interpolating within the task, we can align the latent signals related to the held-out diagonal directions (Fig. S6a, average $r$ over the top five latent dimensions before CCA: -0.05 ± 0.12 (mean ± s.e.); after CCA: 0.44 ± 0.12). As a result, we are still able to use the monkey decoder on the aligned human data, although the average EMG decoding accuracy of the major wrist muscles further decreased to 0.40 compared to when all directions were used (Fig. S6b). On the other hand, when extrapolating beyond the training data, we could no longer align the latent signals of the held-out directions (Fig. S7a, average $r$ over the top five latent dimensions before CCA: 0.17± 0.12 (mean ± s.e.); after CCA: 0.32 ± 0.10). Unsurprisingly, the EMG prediction accuracy for the held-out aligned human latent signals using the source monkey decoder was very low ($R^2 < 0$, Fig. S7b).

**Table S1 | Summary of preferred directions for muscles recorded in common for all three monkeys.**

| | Monkey 1 | | | Monkey 2 | | | Monkey 3 | | |
|---|---|---|---|---|---|---|---|---|---|
| | Session 1 | Session 2 (+29 days) | $\Delta PD$ | Session 1 | Session 2 (+2 days) | $\Delta PD$ | Session 1 | Session 2 (+ 46 days) | $\Delta PD$ |
| ECR | 103° | 103° | 0° | 90° | 96° | 6° | 109° | 129° | 20° |
| EDC | 180° | 180° | 0° | 96° | 109° | 13° | 135° | 154° | 19° |
| FCR | 64° | 58° | 6° | 77° | 71° | 6° | 13° | 6° | 7° |
| FCU | 315° | 321° | 6° | 270° | 283° | 13° | 290° | 283° | 7° |
| FDP | 0° | 0° | 0° | 0° | 347° | 13° | 315° | 341° | 26° |

*The following muscles are evaluated: extensor carpi radialis (ECR), extensor digitorum communis (EDC), flexor carpi radialis (FCR), flexor carpi ulnaris (FCU), flexor digitorum profundus (FDP). Note that additional muscles were recorded for each monkey but not reported here.*

### Supplementary methods

**Behavior tasks**. The isometric box task requires the monkey to control the cursor on the screen by exerting forces on a small box placed around one of the hands. The box was padded to comfortably constrain the monkey's hand and minimize its movement within the box, and the forces were measured by a 6 DOF load cell (JR3 Inc., CA) aligned to the wrist joint. During the task, flexion/extension force moved the cursor right and left respectively, while force along the radial/ulnar deviation axis moved the cursor up and down. Each trial started with the appearance of a center target requiring the monkeys to hold for a random time (0.2 – 1.0 s), after which one of eight possible outer targets selected in a block-randomized fashion appeared, accompanied with an auditory go cue. The monkey was allowed to move the cursor to the target within 2.0 s and hold for 0.8 s to receive a liquid reward. For both decoding and alignment analyses, we only used the data within each single trial using a window starting 0.5 s and ending 1.0 s after force onset.

**iBCI monkey 1 decoders.** We used a Wiener cascade (5) as the fixed source monkey iBCI decoder:

$$\boldsymbol{y}(t) = ReLU\left(\sum_{\tau=0}^{T-1} \boldsymbol{\beta}(\tau)\boldsymbol{x}(t-\tau)\right) \tag{1}$$

where $\boldsymbol{y}(t)$ is a *q*-dimensional vector (*q* varied with the number of recorded EMGs for each monkey) representing the EMGs to be predicted at time *t*, while $\boldsymbol{x}(t)$ is a *p*-dimensional vector for the inputs to the Wiener filter at time *t*, and $\boldsymbol{\beta}(\tau)$ is a $q \times p$ matrix corresponding to the filter parameters for time step $\tau$. $\boldsymbol{x}(t)$ is the projection of the neural firing rates in a linear latent space found by PCA. We set *p* = 13. We can also write (1) in matrix form:

$$\boldsymbol{Y} = ReLU(\boldsymbol{XB}) \tag{2}$$

where $\boldsymbol{Y}$ is a $M \times q$ matrix for the EMGs to be predicted with *M* being the number of samples, $\boldsymbol{X}$ is a $M \times (T \times p)$ matrix, and $\boldsymbol{B}$ is a $(T \times p) \times q$ matrix for the regression coefficients to be estimated. We also added an additional bias term for both $\boldsymbol{X}$ and $\boldsymbol{B}$. $\boldsymbol{B}$ was determined by a ridge regression estimator:

$$\hat{\boldsymbol{B}} = (\boldsymbol{X}^\mathsf{T}\boldsymbol{X} + \lambda\boldsymbol{I})^{-1}\boldsymbol{X}^\mathsf{T}\boldsymbol{Y} \tag{3}$$

We chose a ridge regression to limit the risk of decoder overfitting by penalizing solutions with large regression coefficients with the regularization term $\lambda$. The value of $\lambda$ was chosen by sweeping a range of 20 values between 10 and $10^5$ on a logarithmic scale. We used a 4-fold cross validation to train the fixed decoder for each source monkey and ultimately selected the model with the highest $R^2$ on the test set.

**Supplementary figures**

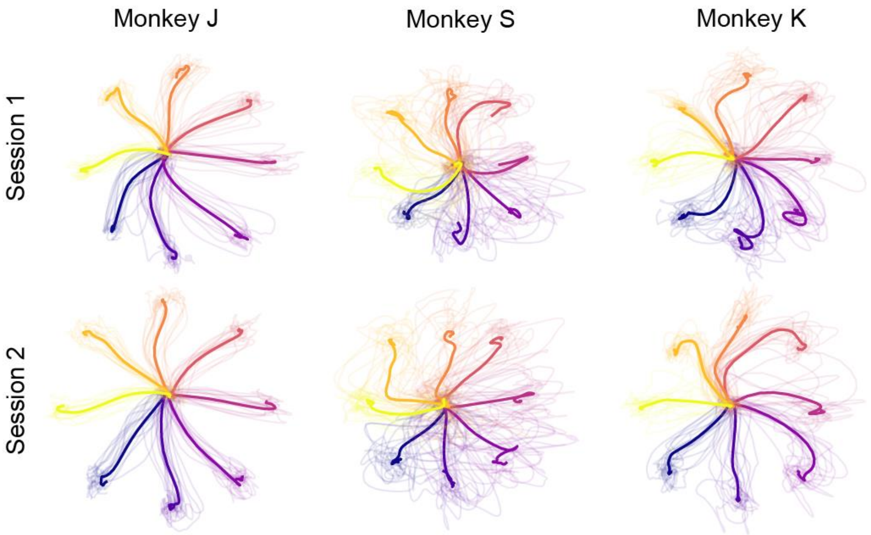


**Figure S1 | Monkeys have slightly different cursor trajectories.** Cursor trajectories of the three monkeys during the two sessions analyzed in this study. Data is averaged across all trials for each target direction (single trial trajectories are shown as shaded curves). Among the three monkeys, Monkey J has the straightest trajectories. Monkey K and (especially) S show the highest trial-by-trial variations.

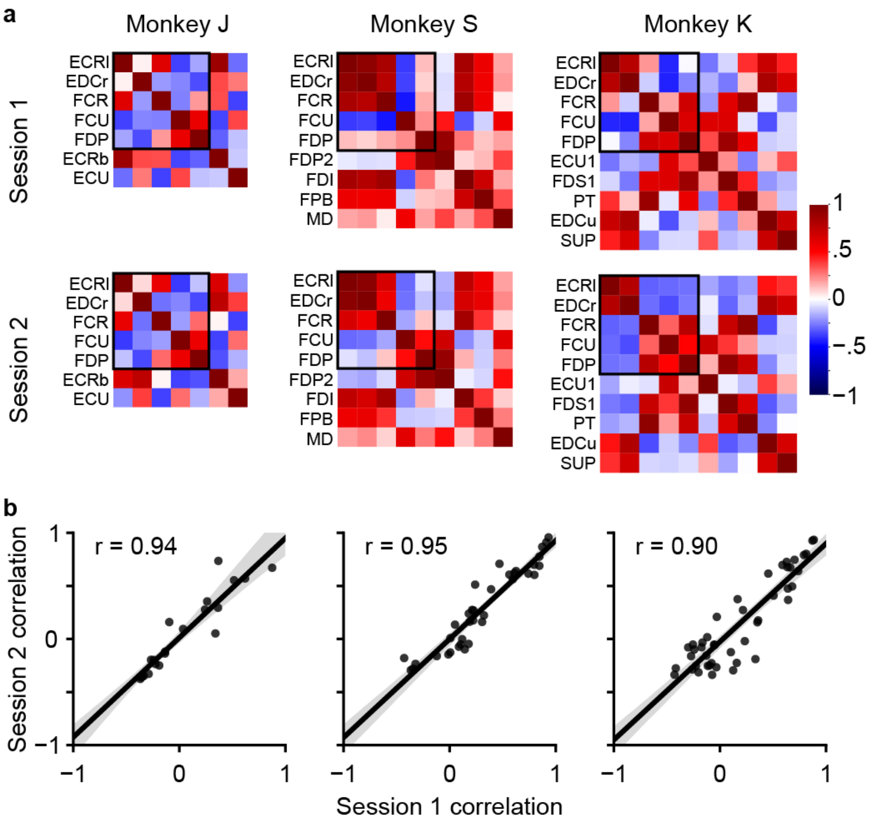


**Figure S2 | Muscle correlation patterns are preserved across time. a**, Matrices of EMG correlations in each monkey during session 1 (top row) and 2 (bottom row). Each matrix is ordered to cluster muscles that have been recorded in common across all three monkeys (indicated by the black square). **b**, Element-by-element scatter plots of the matrices in the top (x-axis) and bottom (y-axis) row of **a.** Correlation of EMGs in the same monkey are very consistent across time.

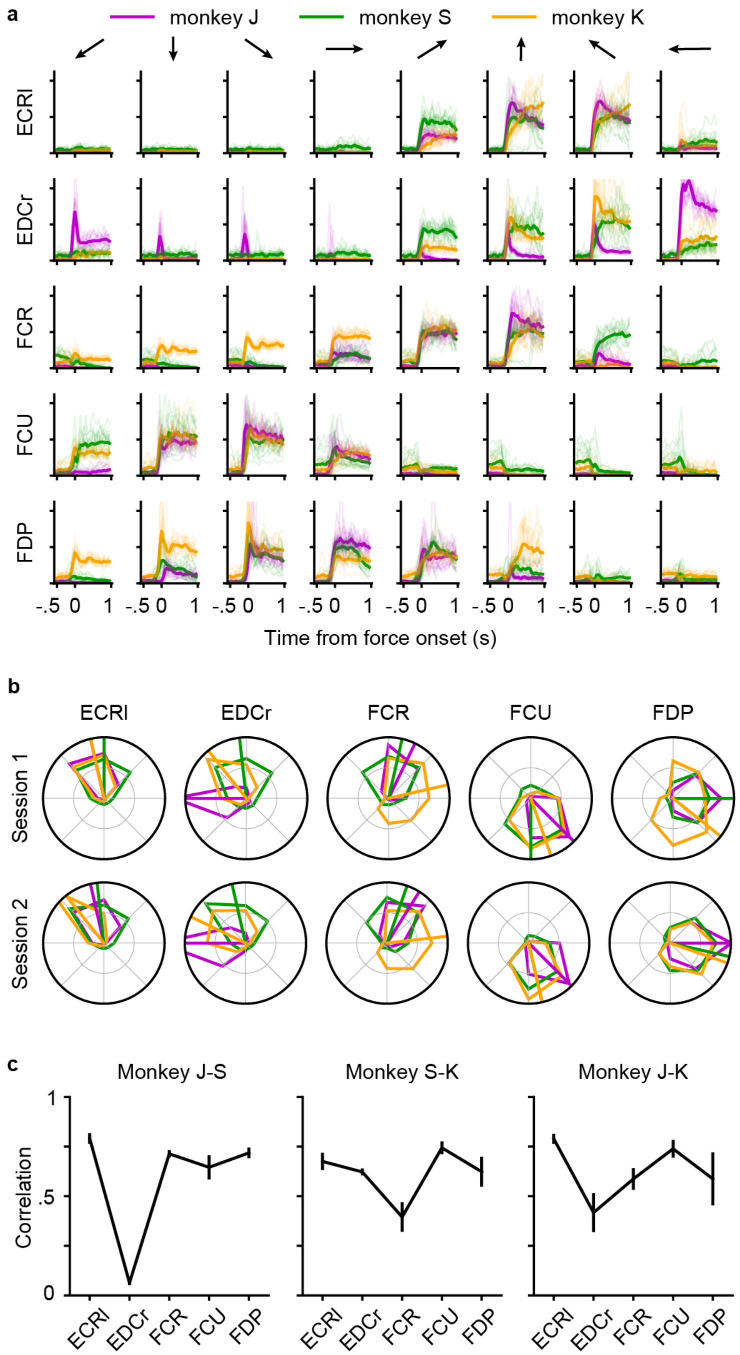


**Figure S3 | Monkeys perform the same task using slightly different muscle strategies**. **a**, EMG traces recorded during the isometric box task for the common muscles of monkey J (purple), monkey S (green), and monkey K (orange). Columns show the average EMG activity for the eight target directions (shaded lines are single trials). The EMG traces are centered around force onset. **b**, Muscle tuning curves expressing the level of activity as a function of the direction of the isometric force. The radial vectors indicate the preferred directions. Top row shows the tuning curves and PDs of monkey J (purple), monkey S (green), and monkey K (orange) recorded during the first session, while bottom row during the second session. **c**, Correlation of EMGs that have been recorded in common between Monkey J and S (left), Monkey S and K (center) and Monkey J and K (right). Mean and standard deviation across all sessions is reported.

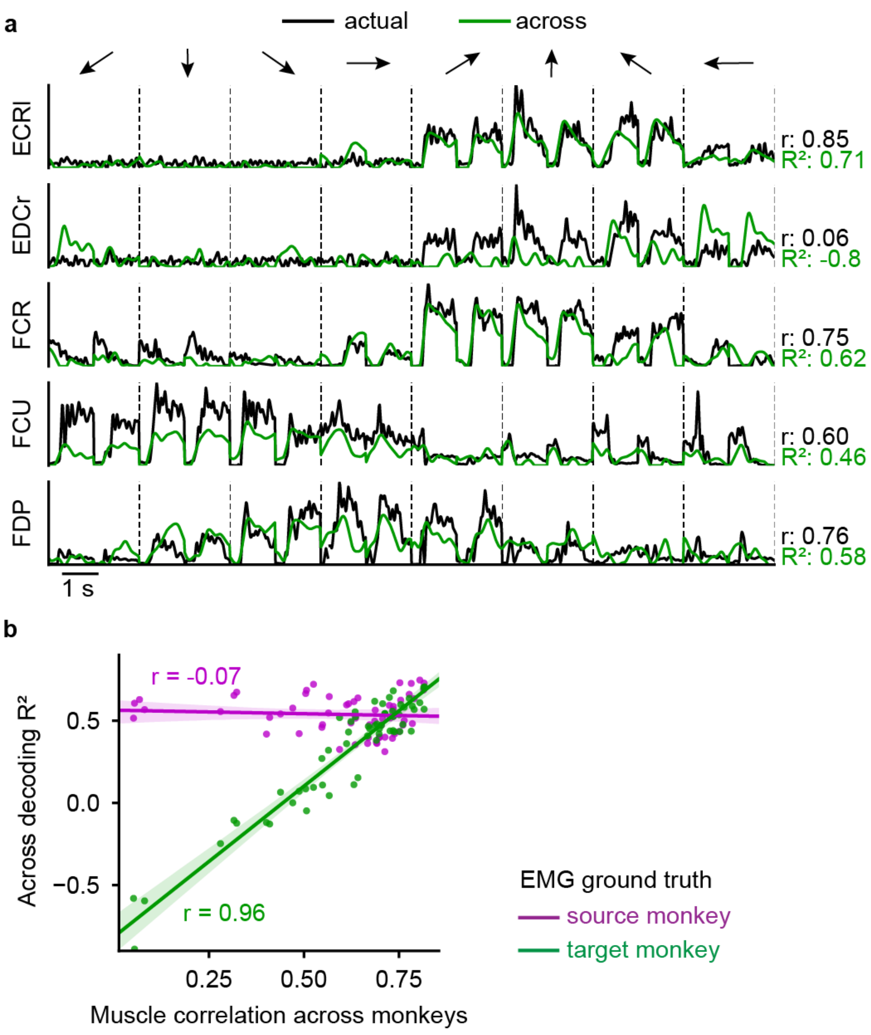


**Figure S4 | Using target monkey EMGs as ground truth for cross-monkey decoding. a**, EMG predictions obtained by feeding the aligned latent signals of the target monkey to the source monkey decoder (green lines) plotted against the actual EMG recordings of the target monkey (black lines). Note that we cannot compare the cross-monkey predictions with the entire set of recorded muscles of the target monkey, but only with the set of EMGs that have been recorded in common across monkeys, as the fixed decoder has been trained to predict the actual EMGs of the source monkey. For each muscle, the cross-monkey predicted $R^2$ is reported with the correlation between the actual EMG recordings of the source and the target monkey. **b**, Muscle correlation between source and target monkeys is plotted against the single muscle cross-monkey $R^2$ when using source (purple) and target (green) monkey EMGs as ground truth. We can decode target monkey EMGs that have high correlation with those of the source monkey. Each dot refers to a single muscle for a given source/target monkey pair.

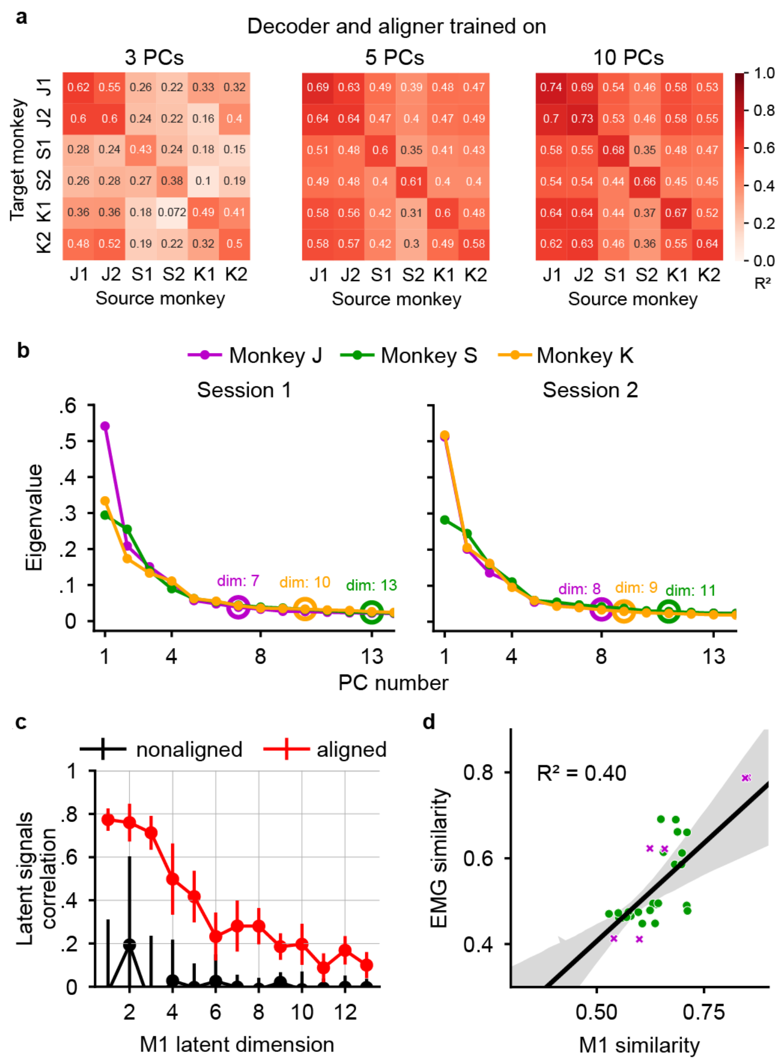


**Figure S5 | Dimensionality analysis. a**, Overall cross-monkey $R^2$ after CCA alignment for all pairs of monkeys shown as a confusion matrix when using different latent space dimensionality. **b**, Scree plot of the first 13 principal components for the first (left) and second (right) session of monkey J (purple), monkey S (green) and monkey K (orange). The Parallel Analysis estimate of linear dimensionality for each monkey's latent signal is indicated as a circle of the corresponding color. **c**, Correlation between the latent signals of source and target monkeys on each of the first 13 latent dimensions before (black) and after (red) CCA alignment. The increased correlation magnitude indicates increased similarity across monkeys as a result of the CCA alignment. The mean and standard deviation across all pairs of monkeys is shown. **d**, M1 similarity is plotted against the EMG similarity across monkeys (green dots) and within monkey/across sessions (purple x symbols). The similarity is computed as the correlation between the CCA-aligned M1/EMG latent signals averaged across the first five latent dimensions. For the EMG similarity, we applied PCA on the EMG signals that were recorded in common across monkeys (see Table S1) and then compute CCA alignment on the corresponding latent signals.

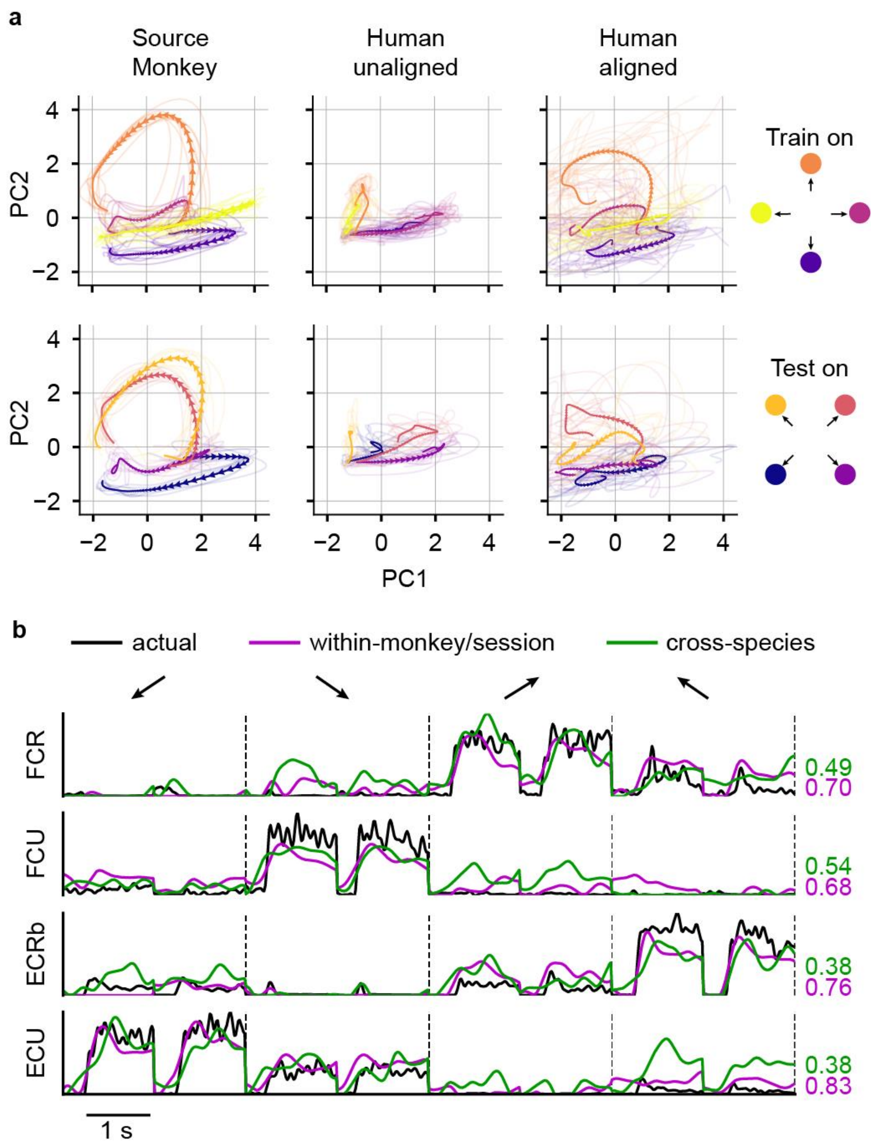


**Figure S6 | CCA alignment for cross-species decoding generalizes when interpolating. a**, Latent M1 trajectories described by the first two principal components of the source monkey (left) and the paralyzed patient attempting to perform a wrist isometric task before (center) and after (right) CCA alignment. The first row shows the latent signals of the directions used to obtain the CCA transform, while the bottom row shows the latent signals of the held-out directions. In both cases, CCA made the neural trajectories of the human similar to those of the source monkey. **b**, Corresponding EMG predictions of the four major wrist muscles on the held-out target directions when feeding the fixed source monkey decoder with source monkey neural data (purple lines) and with the aligned latent signals of the human (green lines). Actual EMG recordings of the source monkey (black lines) are used as ground truth. Vertical dashed lines separate muscle traces for the four held-out target directions.

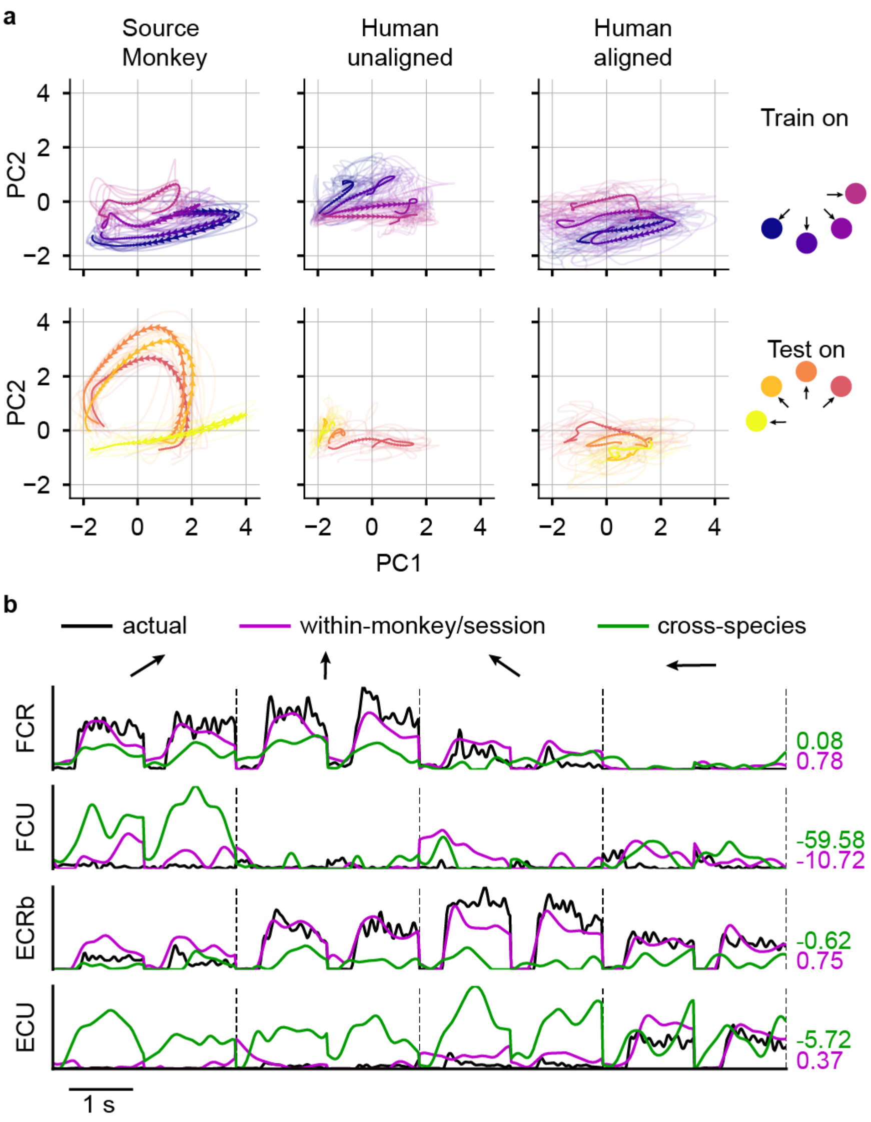


**Figure S7 | CCA alignment for cross-species decoding does not generalize when extrapolating. a**, Latent M1 trajectories described by the first two principal components of the source monkey (left) and the paralyzed patient attempting to perform a wrist isometric task before (center) and after (right) CCA alignment. The first row shows the latent signals of the directions used to obtain the CCA transform, while the bottom row shows the latent signals of the held-out directions. While CCA accurately aligned the latent signals of the training directions, the latent signals of the human bear no resemblance to those of the source monkey in the held-out targets. **b**, Corresponding EMG predictions of the four major wrist muscles on the held-out target directions when feeding the fixed source monkey decoder with source monkey neural data (purple lines) and with the aligned latent signals of the human (green lines). Actual EMG recordings of the source monkey (black lines) are used as ground truth. Vertical dashed lines separate muscle traces for the four held-out target directions.

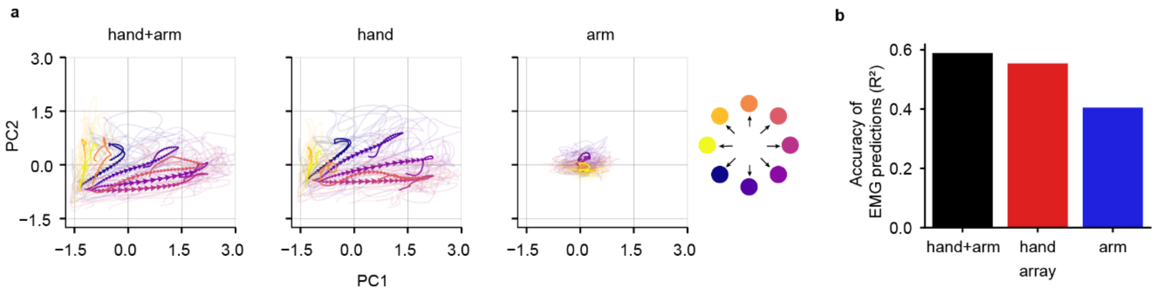


**Figure S8 | Additional cross-species decoding analysis. a**, Latent signals described by the first two principal components of the human recording using channels from both hand and arm (left), only hand (center) and only arm (right) area of M1. Data is averaged across all trials for each target direction (single trial trajectories are shown as shaded curves). Arrows indicate the temporal evolution of the trajectories (0.76 s before to 0.74 s after go cue). **b**, Cross-species $R^2$ when aligning the source monkey latent signals with those resulting from the hand (red), arm (blue) and hand+arm (black) area of the human M1 array. For the cross-species decoding, we only considered the four main wrist muscles.
